# A Single-Cell Atlas of RNA Alternative Splicing in the Glioma-Immune Ecosystem

**DOI:** 10.1101/2025.03.26.645511

**Authors:** Xiao Song, Deanna Tiek, Minghui Lu, Xiaozhou Yu, Runxin Wu, Maya Walker, Qiu He, Derek Sisbarro, Bo Hu, Shi-Yuan Cheng

## Abstract

Single-cell analysis has refined our understanding of cellular heterogeneity in glioma, yet RNA alternative splicing (AS)—a critical layer of transcriptome regulation—remains underexplored at single-cell resolution. Here, we present a pan-glioma single-cell AS analysis in both tumor and immune cells through integrating seven SMART-seq2 datasets of human gliomas. Our analysis reveals lineage-specific AS across glioma cellular states, with the most divergent AS landscapes between mesenchymal- and neuronal-like glioma cells, exemplified by AS in *TCF12* and *PTBP2*. Comparison between core and peripheral glioma cells highlights AS-redox co-regulation of cytoskeleton organization. Further analysis of glioma-infiltrating immune cells reveals potential isoform-level regulation of protein glycosylation in regulatory T cells and a link between *MS4A7* AS in macrophages and clinical response to anti-PD-1 therapy. This study emphasizes the role of AS in glioma cellular heterogeneity, highlighting the importance of an isoform-centric approach to better understand the complex biological processes driving tumorigenesis.

## Introduction

Diffuse gliomas are a heterogeneous collection of devastating brain tumors that vary in cell-of-origin, genetic profile, and clinical behavior^1^. In adult gliomas, the latest WHO classification^2^ is based on *IDH1*/*2* mutations and chromosome (Chr) 1p/19q co-deletion, defining three major subgroups: IDH-mutant (mut), 1p/19q co-deleted oligodendroglioma (IDHO); IDH-mut, 1p/19q-intact astrocytoma (IDHA), both of which are typically lower-grade at diagnosis but inevitably recur as high-grade; and IDH-wildtype (wt) glioblastoma (GBM), a grade IV tumor with an exceptionally poor prognosis, characterized by additional genetic alterations such as Chr 7 gain and Chr 10 loss. Pediatric high-grade gliomas (pHGGs), while sharing a similarly dismal prognosis, have distinct genetic drivers, most notably histone H3 mutations such as H3-K27M and H3-G34R/V. Although less common, some pediatric gliomas also harbor IDH mutations, suggesting a potential overlap in tumorigenic mechanisms between certain adult and pediatric gliomas.

Single-cell RNA-sequencing (scRNA-seq) has revolutionized our understanding of cellular heterogeneity in glioma. Both adult and pediatric glioma cells hijack early neural developmental programs under genetic and environmental drivers, leading to heterogenous and plastic transcriptomic states that resemble neurodevelopmental lineages, albeit in a partial or distorted form^3–8^. Additionally, the immune compartment in glioma tumor microenvironment (TME), particularly myeloid cells and T cells, has also been extensively investigated through scRNA-seq^9–13^. These studies have revealed immunosuppressive and tumor-promoting properties of distinct myeloid cell populations, as well as mechanisms underlying impaired T cell function in gliomas, highlighting novel therapeutic opportunities by rewiring immune cell states.

Alternative splicing (AS), a prevalent process that enables a single gene to produce multiple mRNA isoforms, is an important domain of transcriptomic heterogeneity^14^. In gliomas, AS dysregulation has been largely studied through bulk-level analyses^15–19^, which are significantly influenced by tumor purity of the bulk tissues. Accumulated scRNA-seq studies in gliomas have solely focused on overall gene expression levels to classify cell subpopulations or identify potential therapeutic targets, while the isoform-level alterations have largely been overlooked. It remains unclear whether glioma tumor cells exploit AS as a gene regulatory mechanism to accommodate transitions among different cellular states, and whether immune cells express distinct isoforms to adapt their functions in glioma TME. To address these gaps, we performed a pan-glioma single-cell AS analysis by integrating seven published SMART-Seq2 datasets from clinical glioma samples^4–6,8,12,20,21^, along with scRNA-seq data from non-tumor brains and peripheral blood mononuclear cell (PBMC) samples^22,23^. These SMART-Seq2 datasets provide full mRNA coverage, unlike droplet-based scRNA-seq that captures only the 5’ or 3’ ends of the mRNA, making them ideal for AS estimation. Our analysis revealed a distinct AS landscape in mesenchymal-like glioma cells compared with neuronal-like glioma cells. Further analysis of GBM tumor cells in the core and peripheral areas highlighted the significant roles of AS in regulating actin organization. Additionally, analysis of glioma-infiltrating immune cells reveals potential isoform-level regulation of protein glycosylation in regulatory T cells and distinct AS landscapes between microglia and blood-derived monocytes/macrophages. Overall, our analysis provides a single-cell AS atlas of both tumor and immune cells in human gliomas, offering a promising framework for future research and therapeutic development.

## Results

### The lineage preference of adult and pediatric glioma cells is associated with genetic background and spatial distribution within the tumor

We compiled a SMART-Seq2 single-cell transcriptome atlas in 36 adult GBMs, 31 adult IDH-mut gliomas, ten H3-WT pHGG, six K27M pHGG, nine G34R pHGG, 12 non-tumor brains and pooled non-tumor PBMC samples from previously published datasets^4–6,8,12,20–23^(Tables S1-2). Uniform manifold approximation and projection (UMAP)-based clustering of 32,304 high-quality cells generated a total of 29 clusters (Fig. S1A-B and Tables S3-4). Non-tumor cell types were assigned based on canonical markers (Fig. S1B), including T cell (*CD2*, *CD3E*), B cell (*CD19*, *CD79A*), macrophage/microglial/monocyte (*CD14*, *CSF1R*), neutrophil (*CSF3R*, *FCGR3B*), astrocyte (*AQP4*, *ADGRV1*), neuron (*RBFOX1*, *NRXN3*), oligodendrocyte (*MBP*, *MOBP*), fibroblast (*COL1A1*, *COL3A1*), and choroid plexus (*PTPRQ*^24^, *CFTR*^25^). Malignant cells were identified based on inferred copy number alterations using copyKAT^26^ (Fig. 1C and S1C-H). This analysis identified large-scale amplifications and deletions in predicted aneuploid cells, including Chr 7 gain and Chr 10 loss in adult GBM cells, and 1p/19q co-deletion in IDHO samples. Single-nucleotide variant calling in *IDH1* and *H3* genes further validated the aneuploid prediction by copyKAT (Fig. 1D and S1C-H).

**Figure 1.**
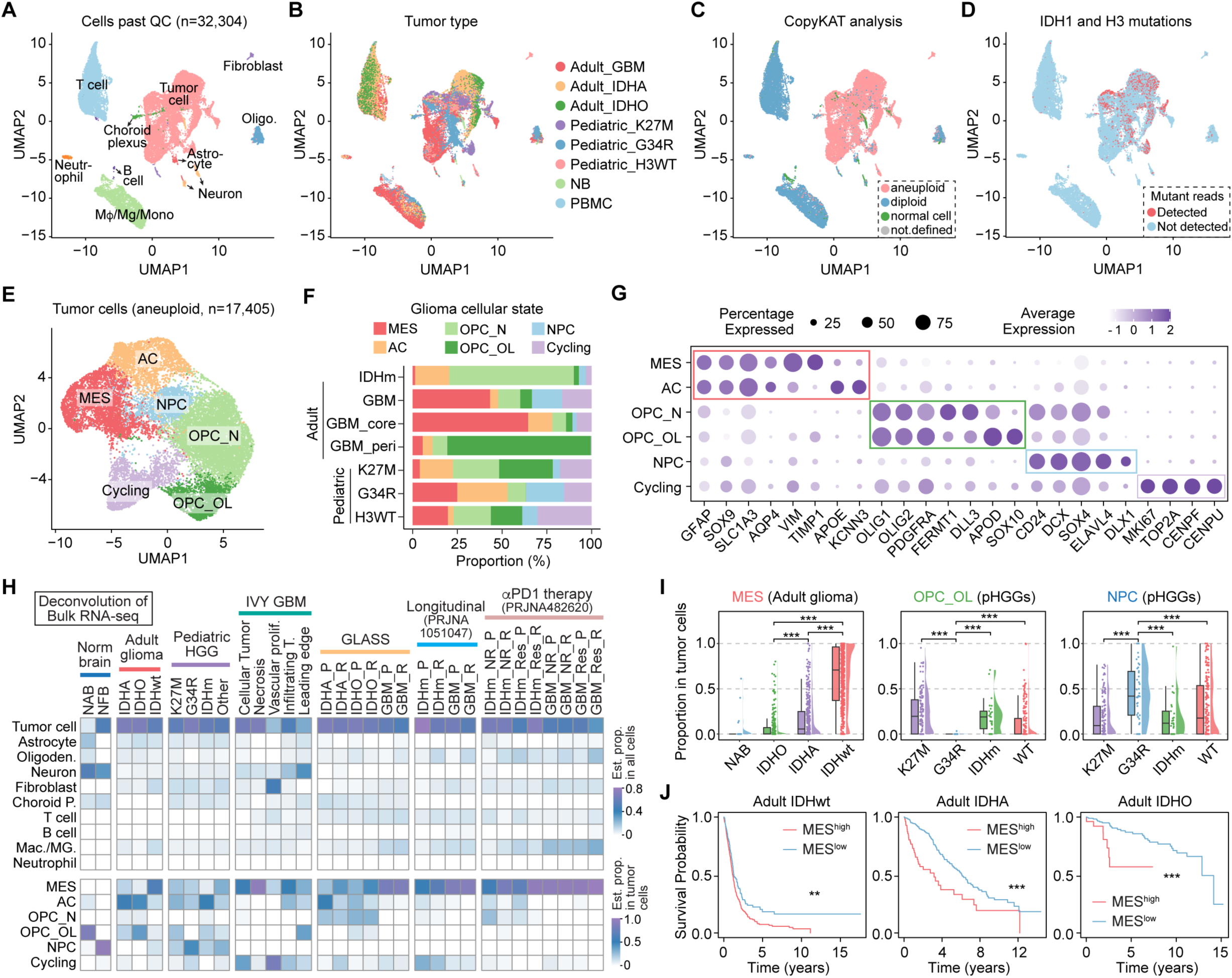
The lineage preference of adult and pediatric glioma cells is associated with genetic background and spatial distribution within the tumor. **A-D**. UMAP projections of all high-quality cells colored by cell type assignment (**A**), sample type (**B**), diploid/aneuploid status determined by CopyKAT (**C**), and detection of *IDH1*/*H3* mutant reads (**D**). **E.** UMAP projections of malignant cells colored by assigned cellular state. **F.** Bar graph summarizing the compositions of glioma cellular states from each glioma subtype. **G.** Dot plot showing marker gene expression. **H.** Deconvolution in glioma bulk RNA-seq data showing the predicted compositions of malignant and non-malignant cell types. NAB, normal adult brain. NFB, normal fetal brain. P, primary. R, recurrent. Res, responder. NR, non-responder. **I.** Raincloud plot showing the proportions of MES, OPC_OL, and NPC glioma cellular states across glioma subtypes, as predicted by deconvolution analysis. **J.** Kaplan-Meier analyses of overall survival across adult glioma subtypes, stratified by the predicted proportion of the MES cells in all tumor cells. The cutoff of MES proportion is 0.4 (IDHwt), 0.35 (IDHA), or 0.2 (IDHO). **, p < 0.01. ***, p < 0.001.

Within malignant cells, UMAP-based clustering identified six major glioma cellular states (Fig. 1E-G and S2A-B). A mesenchymal-like (MES) state is significantly enriched in adult GBMs, particularly in samples from the tumor core (Fig. 1F). In contrast, peripheral GBM cells predominantly exist in a cellular state characterized by high expression of both oligodendrocyte progenitor cell (OPC) markers (*OLIG1*, *OLIG2*, *PDGFRA*) and pre-mature oligodendrocyte (OL) markers (*APOD*, *SOX10*), defining the OPC_OL state. Adult IDH-mut tumor cells exhibit fewer cycling cells (*MKI67*, *TOP2A*), with the majority highly expressing OPC markers while also showing intermediate expression of neuronal progenitor cell (NPC) markers (*CD24*, *DCX*, *SOX4*), defining a state termed OPC_N. Among pHGGs, the G34R subtype is devoid of the OPC_OL state and instead shows an increasing proportion of NPC-like states, characterized by high expression of NPC markers (Fig. 1G).

To validate these differential lineage preferences across glioma subtypes, we performed a deconvolution analysis in several bulk glioma RNA-seq datasets^27–34^ (Table S5). The results confirmed the enrichment of MES state around necrotic regions in adult GBMs (Fig. 1H-I). Additionally, the proportion of MES cells significantly correlated with a worse prognosis in all adult glioma subtypes (Fig. 1J). In pHGGs, G34R tumors lack the OPC lineages and instead have the highest proportion of NPCs among pHGG subtypes (Fig. 1I), further validating the scRNA-seq analysis. In conclusion, both scRNA-seq and bulk RNA-seq deconvolution analyses provide complementary evidence for the lineage preferences in glioma cells, which are linked to genetic background and spatial distribution within the tumor.

### Single-cell AS analysis in glioma tumor cells reveals lineage-specific AS regulation in *PTBP2* and *TCF12*

To interrogate the AS heterogeneity within glioma cells, we quantified the Percent Spliced In (PSI) value for each detected AS event in individual cell using the MARVEL package^35^. MARVEL detected seven types of events: SE (skipped exon), MXE (mutually exclusive exons), A5SS/A3SS (alternative 5’/3’ splice sites), AFE/ALE (alternative first/last exons), and RI (intron retention). We excluded RI from our analysis, as it usually requires higher sequencing depth for accurate quantification^36^. Additionally, some of the included datasets contain single-nucleus RNA-seq data that captures unsliced transcripts, further limiting the RI detection. Next, we identified 2,531 variable AS events, defined as those with a standard deviation (SD) of PSI > 0.2 and detected in more than 500 cells (Fig. S3A and Table S6). SE is the most prevalent type among these events. The UMAP projection based on the PSI data of these 2,531 events revealed a separation of IDH-mut tumor cells from others, indicating distinct AS landscape associated with IDH mutation (Fig. 2A). Notably, MES states were distant from OPC-N and a small subset of NPC in the UMAP plot, indicating the greatest AS divergence among these states. This is further supported by the largest number of differentially spliced events observed between MES and OPC-N among all state pairs (Fig. 2B and Table S7). Moreover, most of the differentially spliced events between MES and OPC-N show less than a two-fold difference in overall gene-level expression (Fig. S3B), suggesting that AS is a separate mechanism in regulating gene expression across glioma cellular states, where these differences would be lost in a global gene expression analysis.

**Figure 2.**
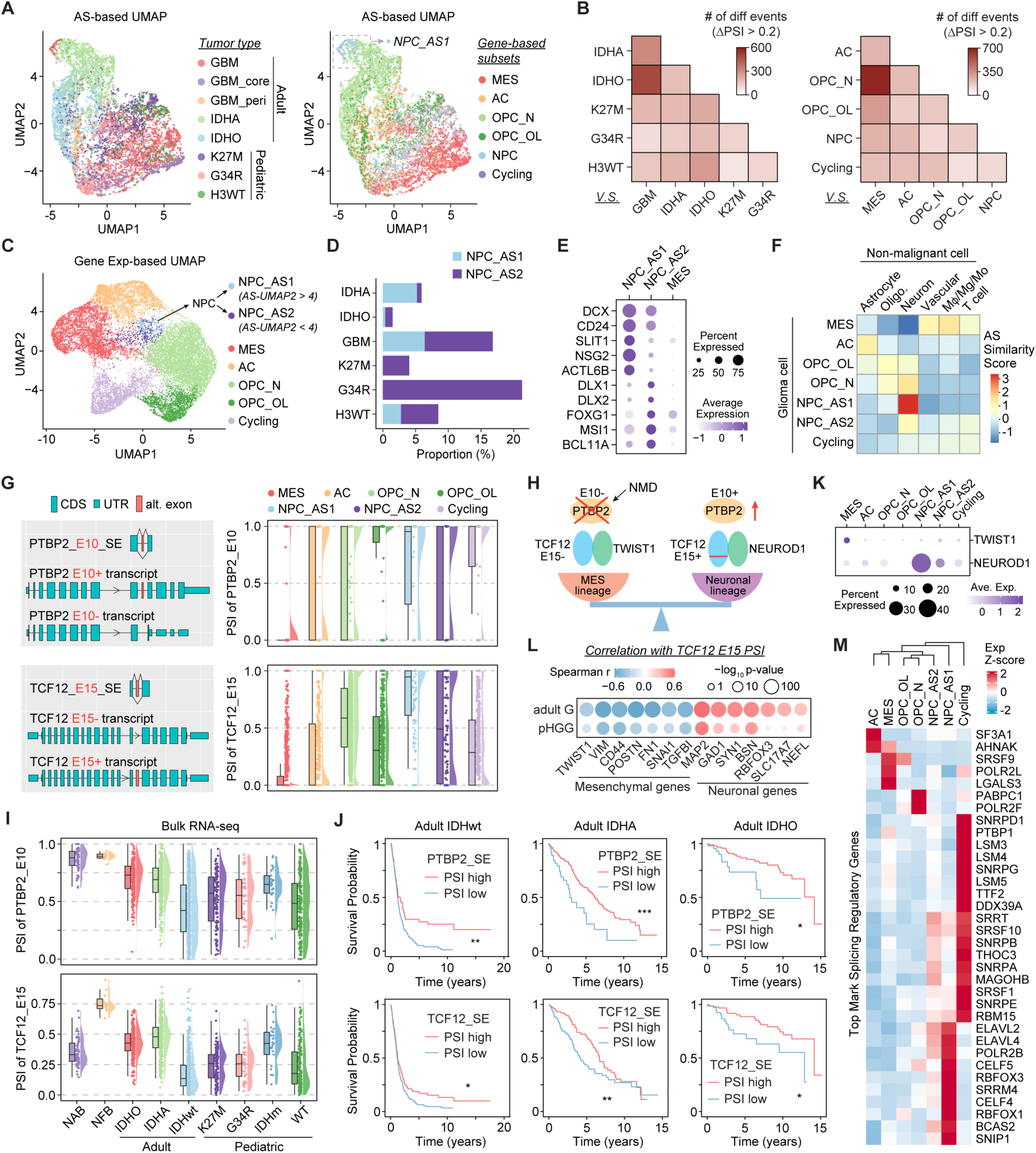
Single-cell AS analysis in glioma tumor cells reveals lineage-specific AS regulation in *PTBP2* and *TCF12*. **A**. UMAP projections of glioma cells based on PSI data of variable events. Cells are colored by tumor type (left) or gene-based cellular state (right). **B.** Heatmap showing the number of differential events for each pairwise comparison of glioma subtypes (left) or cellular states (right). **C.** UMAP projections of glioma cells based on the gene expression landscapes. Cells are colored by gene-based cellular state while the cells of NPC subset are divided into NPC_AS1 and NPC_AS2 based on the AS-based UMAP projection in (**A**). **D.** Bar graph summarizing the compositions of NPC_AS1 and NPC_AS2 from each glioma subtype. **E.** Dot plot showing marker gene expression for NPC_AS1 and NPC_AS2, with MES as a control. **F.** Heatmap showing the AS similarity between each pairwise comparison of non-malignant cell types and glioma cellular states. **G.** Diagrams (left) illustrating the exon structures of the indicated AS events, and raincloud plots (right) showing the PSI distribution of these AS events across glioma cellular state. **H.** Schematic illustrating the hypothesis that AS in *PTBP2* and *TCF12* contribute to cell state transitions between neuronal and mesenchymal lineages. **I.** Raincloud plots showing the PSI distribution of the indicated events across glioma subtypes and normal controls from bulk RNA-seq data. **J.** Kaplan-Meier analyses of overall survival across adult glioma subtypes, stratified by the PSI values of indicated events. **K.** Dot plots showing the expression of *TWIST* and *NEUROD1* across glioma states. **L.** Dot plot showing the correlation between *TCF12*_E15 PSI and the expression of genes associated with mesenchymal and neuronal lineages. **M.** Heatmap showing the gene expression of splicing factors across glioma cellular states. *, p < 0.05. **, p < 0.01. ***, p < 0.001.

AS-based UMAP analysis revealed two distinct clusters within the NPC state, a separation not observed in gene-based UMAP (Fig. 2C). We designated NPC state cells with AS-UMAP2 > 4 as the NPC_AS1 state, and those with AS-UMAP2 < 4 as the NPC_AS2 state. The proportion of these two NPC states varies among glioma subtypes, with the G34R subtype exhibiting the highest NPC_AS2 proportion (Fig. 2D). NPC_AS1 cells express higher levels of *DCX*, *CD24*, and mature neuron markers such as *SLIT1*^37^ and *NSG2*^38^. Conversely, NPC_AS2 cells preferentially express interneuron markers, including *DLX1*, *DLX2*, and *FOXG1* (Fig. 2E). This finding aligns with previous studies indicating an interneuronal lineage as the putative origin of G34R tumors^6,31^.

To determine whether the differential AS landscapes among glioma states result from lineage-specific AS, we focused on the top 631 differential events (ΔPSI > 0.3) across glioma states and compared the similarity of their PSI values between glioma cells and non-malignant cells. Surprisingly, NPC_AS1 cells exhibit AS patterns closely resembling those of neurons, whereas the MES state shows the least similarity to the three neural lineages and aligns more closely with non-neural cells, particularly the myeloid lineage, indicating their mesodermal lineage commitment (Fig. 2F). The AC, OPC_OL, OPC_N, and NPC_AS2 states display AS landscapes that are slightly closer to their corresponding normal lineage cells: astrocytes, oligodendrocytes, or neurons. These findings suggest that AS landscape in glioma tumor cells reflects their lineage states.

Next, we focused on two SE events among the top differential events between MES and OPC_N states (Table S7): *PTBP2*_E10_SE and *TCF12*_E15_SE, occurring in genes that are important regulators of neuronal or mesenchymal lineage gene expression^39–42^. Polypyrimidine tract-binding proteins 1 and 2 (PTBP1 and PTBP2) are well-characterized splicing repressors^43^. PTBP1, ubiquitously expressed across various tissues, is a strong repressor of neuron-specific exons. Conversely, PTBP2, predominantly expressed in neuronal cells, demonstrates a much less repressive capacity on neuronal exons. The transition from PTBP1 to PTBP2 expression is a crucial event during neuronal differentiation, facilitating the inclusion of neuronal exons essential for proper neuronal maturation^39^. The MES glioma cells predominantly express E10-excluded *PTBP2* isoform (Fig. 2G and S3D). This SE event introduces a premature stop codon that triggers NMD, leading to reduced protein production^44^. The decreased *PTBP2* expression may contribute to the reduced neuronal lineage commitment observed in MES cells (Fig. 2H). As expected, decreased inclusion of *PTBP2*-E10 was observed in the bulk RNA-seq data of IDHwt gliomas compared with IDH-mut gliomas and normal brains (Fig. 2I), and this decreased inclusion was associated with a worse prognosis (Fig. 2J).

TCF12 is a member of the basic helix-loop-helix (bHLH) transcription factors, which function by forming homodimers or heterodimers with other bHLH proteins to regulate gene transcription during development^45^. The TCF12-TWIST1 dimers drive the mesenchymal transition^40,41^, while the TCF12-NEUROD1 dimer is essential for the induction of neuronal genes during cortical development^42^. The E15-excluded *TCF12* is the dominant isoform in MES states, whereas the E15-included isoform is preferentially expressed in the neuronal lineages, including NPC_AS1 and OPC-N (Fig. 2G). E15 of *TCF12* encodes an in-frame ankyrin-like domain of 24 amino acids (aa) that is inserted into the activation domain (Fig. S3C), which has been reported to affect the DNA binding capacity of TCF12 and its dimer formation^46^. Interestingly, glioma cells in MES and NPC_AS1 states express *TWIST1* or *NEUROD1*, respectively (Fig. 2K), suggesting that TCF12 E15- and E15+ isoforms may regulate mesenchymal and neuronal transcription through forming dimers with TWIST1 and NEUROD1, respectively (Fig. 2H). This hypothesis is supported by the negative correlation between *TCF12*-E15 PSI and the expression of mesenchymal genes, as well as the positive correlation between *TCF12*-E15 PSI and the expression of neuronal genes in glioma bulk RNA-seq data (Fig. 2L). Furthermore, decreased inclusion of *TCF12*-E15 was observed in adult GBMs compared to IDH-mut gliomas, and lower PSI was associated with a worse prognosis (Fig. 2I-J).

AS is tightly regulated by splicing factors (SFs), which determine the precise inclusion or exclusion of exons in mRNA. Differential expression analysis across glioma states identified a group of SFs that are upregulated in cycling cells, including *PTBP1*, *SRSF1*, and *SNRPB* (Fig. 2M), which were previously shown to be upregulated in glioma and promote glioma progression^15,47,48^. In contrast, the SFs upregulated in NPC_AS1 are mostly neuronal-specific SFs^49^. It was reported that PTBP1 represses the inclusion of *PTBP2*-E10^44^, and this was confirmed in our RNA-seq data from patient-derived glioma stem-like cells (GSC1478) treated with shRNA-PTBP1 or shRNA-control (Fig S3D). Surprisingly, we also observed increased inclusion of *TCF12*-E15 following PTBP1 knockdown, suggesting that PTBP1 plays a role in suppressing neuronal-specific exons in glioma cells.

### Comparison between core and peripheral GBM cells highlights AS regulation of cytoskeleton organization

Glioma cells diffusively invade normal brain tissue, contributing to the inevitable recurrence^50^. Therefore, we decided to investigate the transcriptomic regulation in peripheral GBM cells to identify potential therapeutic targets specifically for these cells. Gene-level expression analysis identified differentially expressed genes, including splicing factors, and their enriched pathways in peripheral and core tumor cells (Fig. 3A-B). The following AS analysis identified 173 differentially spliced events between peripheral and core GBM cells (Fig. 3C and Table S8). Interestingly, these affected genes are enriched in pathways related to actin cytoskeleton organization (Fig. 3D). Among the 18 genes involved, three belong to the tropomyosin family (*TPM1-3*), which encode α-helical coiled-coil proteins that stabilize actin filaments by wrapping around them^51^, and one belongs to the α-actinin family (*ACTN4*), which binds and crosslinks actin filaments^52^ (Fig. 3E and S4A).

**Figure 3.**
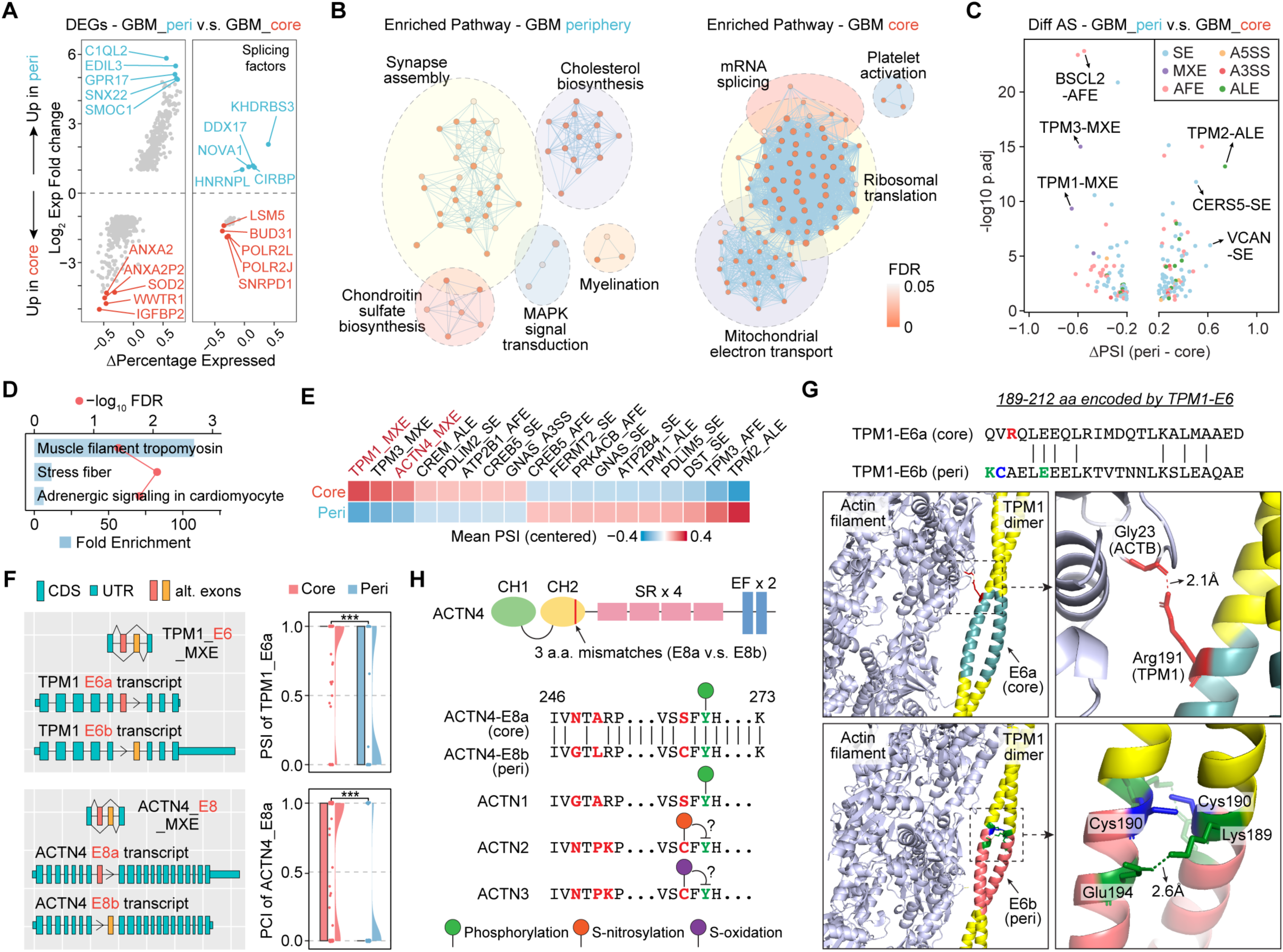
Comparison between core and peripheral GBM cells highlights AS regulation of cytoskeleton organization. **A**. Scatter plot showing the log_2_ fold change of expression (y-axis) and the difference in the percentage of cells with detectable expression of specific gene (x-axis) for differentially expressed genes (left) or splicing factors (right) between peripheral and core GBM tumor cells. **B.** Pathway enrichment analysis of genes with higher expression in peripheral or core GBM cells. **C.** Volcano plot showing differentially spliced events between peripheral and core GBM cells. **D.** Enriched GO biological processes of genes that are differentially spliced between peripheral and core GBM cells. **E.** Heatmap showing the mean PSI of differential evens in peripheral and core GBM cells. **F.** Diagrams (left) illustrating the exon structures of the indicated AS events, and raincloud plots (right) showing the PSI distribution of these AS events in peripheral and core GBM cells. **G.** AlphaFold 3 prediction of the interaction between TPM1 dimer and actin filament, focusing on the region encoded by the mutually exclusive E6. **H.** Sequence comparison of ACTN4 isoforms and reported PTMs. CH, calponin homology actin-binding domain. SR, spectrin repeats, EF, EF-hand motif. ***, p < 0.001.

TPM1 has two mutually exclusive E6, both encoding 189-212 aa, with core GBM cells primarily using the upstream E6a and peripheral cells favoring E6b, yet overall gene expression remains unchanged (Fig. 3F and S4B). TPM1 proteins form homo-dimers that lie along the α-helical groove of actin filaments. AlphaFold 3^53^ predicts that Arg191, encoded by E6a, forms a close bond with Gly23 of ACTB (2.1Å), while E6b encodes a polypeptide with a Cys190 site, which is reported to form a disulfide bridge within the TPM1 dimer, thereby impairing the actin-tropomyosin interaction^54^ (Fig. 3G). Additionally, predicted bonds between two pairs of Lys189 and Glu194 in the TPM1 E6b dimer may further strengthen the disulfide bridge. These findings suggest that peripheral GBM cells express TPM1 E6b isoform with reduced actin binding affinity, particularly under oxidative stress. This may alleviate tropomyosin’s stabilizing and stiffening effects on actin filaments, promoting a more dynamic actin organization that facilitates cell invasion.

Core and peripheral GBM cells also differentially express ACTN4 isoforms with two mutually exclusive E8, leading to three mismatches in the second calponin homology (CH) actin-binding domain (Fig. 3F and 3H). Core GBM cells express the E8a isoform with Asn248, Ala250, and Ser263, while the peripheral isoform with E8b encodes Gly248, Leu250, and Cys263. Notably, the nearby Tyr265 is frequently phosphorylated, according to the PhosphoSite database (Fig. S4C), and its phosphorylation significantly enhances the acting-binding activity^55,56^. Moreover, this tyrosine site is conserved among four α-actinin family members, with phosphorylation also commonly detected in ACTN1, which has a serine at the second upstream position, same as the ACTN4-E8a isoform (Fig. 3H and S4D). In contrast, phosphorylation is rarely observed in the same tyrosine sites of ACTN2/3, which have cysteines at the second upstream positions, same as the ACTN4-E8b isoform. These cysteines in ACTN2/3 have been reported to undergo oxidation or nitrosylation^57,58^, according to the CysModDB database. These findings suggest that the Cys263 substitution in the ACTN4 E8b isoform, expressed in peripheral GBM cells, may modulate Tyr265 phosphorylation as a redox switch, potentially unfreezing the cross-linked actin filaments in peripheral tumor cells.

### Glioma-infiltrating T cells have an increased proportion of Treg subsets and a decreased proportion of NKT subsets

To investigate the transcriptomic heterogeneity of T cells in gliomas, we extracted 7,064 cells from clusters 1, 3, 11, 13, and 29 (Fig. S1A) and performed de novo clustering with gene expression data (Fig. 4A-B). Based on canonical immune markers, differentially expressed genes, and curated gene signatures^59^ (Fig. 4C-D and Table S9), we defined seven T cell subsets: CD4_naive (*CCR7*, *LEF1*), CD4_act (activated, *CD40LG*), CD4_ISG (type I interferon-stimulated genes, *ITIFs*), CD4_Treg (*FOXP3*, *IL2RA*), CD8_GZMK (granzyme K), CD8_NKT (co-expressing *CD3E* and NK marker, *KLRD1*), T_stress (expressing stress-related heat shock genes), and T_oxphos (enriched for oxidative phosphorylation genes). Additionally, we identified an NK cell cluster, characterized by low *CD3E* expression and high level of NK markers (*KLRF1*, *FGFBP2*). For the subsequent analysis, we focused on the seven T cell subsets.

**Figure 4.**
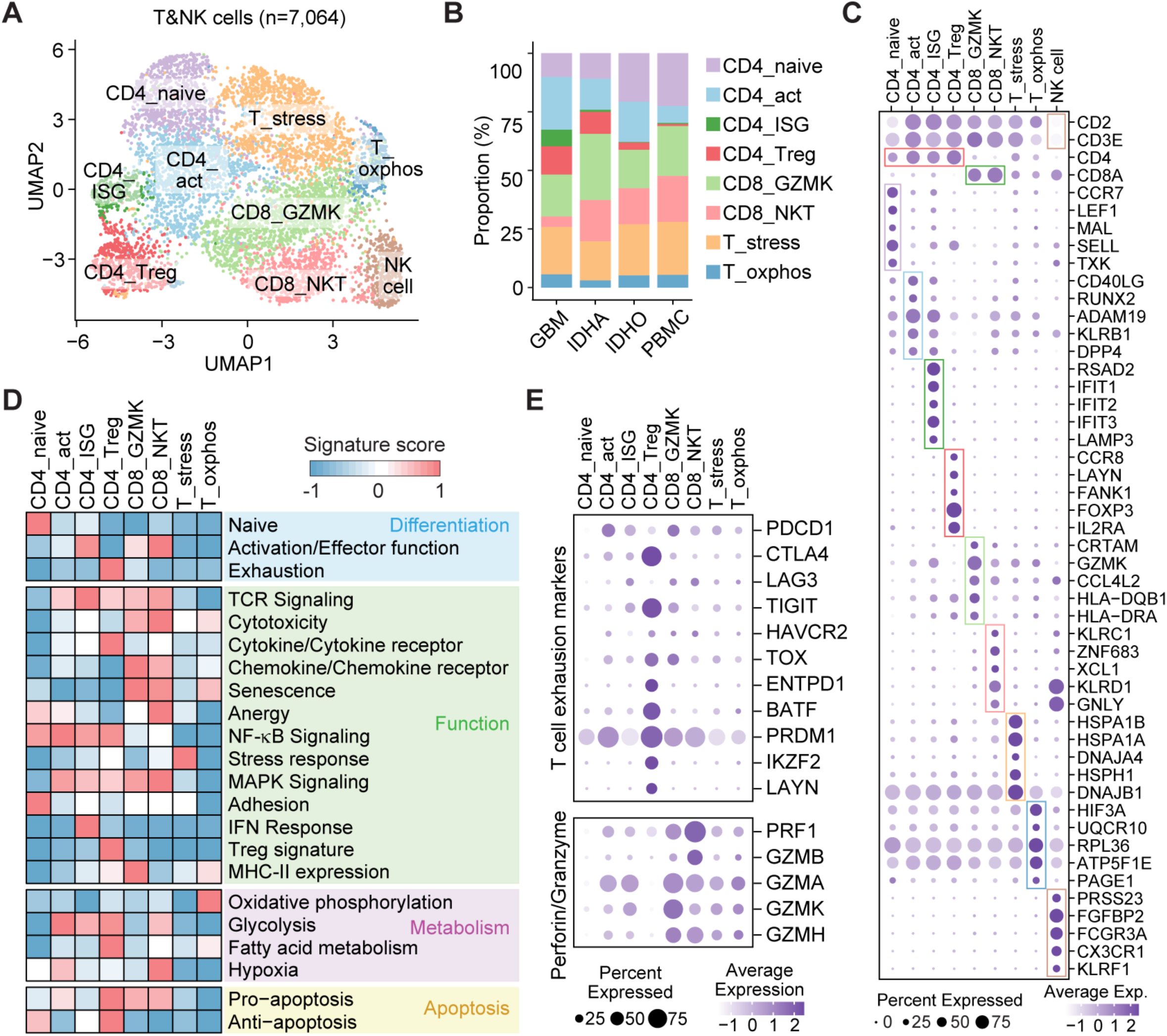
Glioma-infiltrating T cells have an increased proportion of Treg subsets and a decreased proportion of NKT subsets. A. UMAP projections of glioma-associated T and NK cells colored by assigned cellular state. B. Bar graph summarizing the compositions of T cell subsets from indicated groups. C. Dot plot showing marker gene expression. D. Heatmap showing scaled gene signature scores of curated gene signatures across T cell subsets. E. Dot plot showing gene expression across T cell subsets.

T cell exhaustion, a hallmark of chronic antigen exposure, leads to a progressive T cell dysfunction, including impaired cytolytic activity^60^. Analysis of T cell exhaustion markers revealed their highest expression in CD4_Treg (Fig. 4E). Additionally, two CD8+ T cell subsets exhibited differential expression of cytolytic granzyme molecules. Both *PRF1* (perforin) and *GZMB*, which possess strongest cytolytic potential^61^, were expressed at higher levels in the CD8_NKT subset compared to the CD8_GZMK subset. Notably, GBM-infiltrating T cells exhibited a higher proportion of CD4_Treg subsets and a reduced proportion of CD8_NKT cells compared to IDH-mut tumors and normal PBMC controls (Fig. 4B), indicating more pronounced T cell dysfunction in GBM.

### AS regulation in glioma-infiltrating T cells affects genes involved in nucleotide sugar metabolism and calcium entry

The differential AS analysis among T cell subsets revealed much less heterogeneity compared to tumor populations, with a total of 37 events differentially spliced between at least one pair of T cell subsets (Fig. S5A and Table S10). SE remained the most prevalent type among the differential events, followed by AFE (Fig. S5B). Additionally, greater AS heterogeneity was observed in CD4^+^ T cell subsets compared to CD8^+^ T cell subsets (Fig. S5A). One of the events that was differentially spliced in the CD4_Treg population, compared with other CD4^+^ T subsets, is the AFE of UDP-glucose pyrophosphorylase 2 (UGP2, Fig. 5A). CD4_Treg preferentially use the downstream E1b as the first exon, generating an isoform with an additional 11 aa at the N-terminal compared with E1a isoform (Fig. 5B-C). Although the functional impact of these 11 aa remains unclear, several post-translational modifications (PTMs) have been identified in this region, including the methylation of Arg3, which has been detected in primary T cells^62^. UGP2 is the only known enzyme in mammalian cells that catalyzes the reaction of UTP + glucose 1-phosphate → UDP-glucose + PPi^63^. UDP-glucose, the product of this reaction, is the major glucosyl donor for glycogenesis in animals, as well as for the synthesis of various glycoconjugates, such as glycosphingolipid, glycosaminoglycan, and proteoglycans^64^. Interestingly, we observed that CD4+ Tregs not only preferentially express the longer UGP2 isoform but also upregulate the overall expression of UGP2 and several downstream enzymes in the UDP-glucose pathway, particularly those involved in N- and O-glycosylation (Fig. 5D, 5E, and S5C), suggesting a role for protein glycosylation in glioma-associated Treg population.

**Figure 5.**
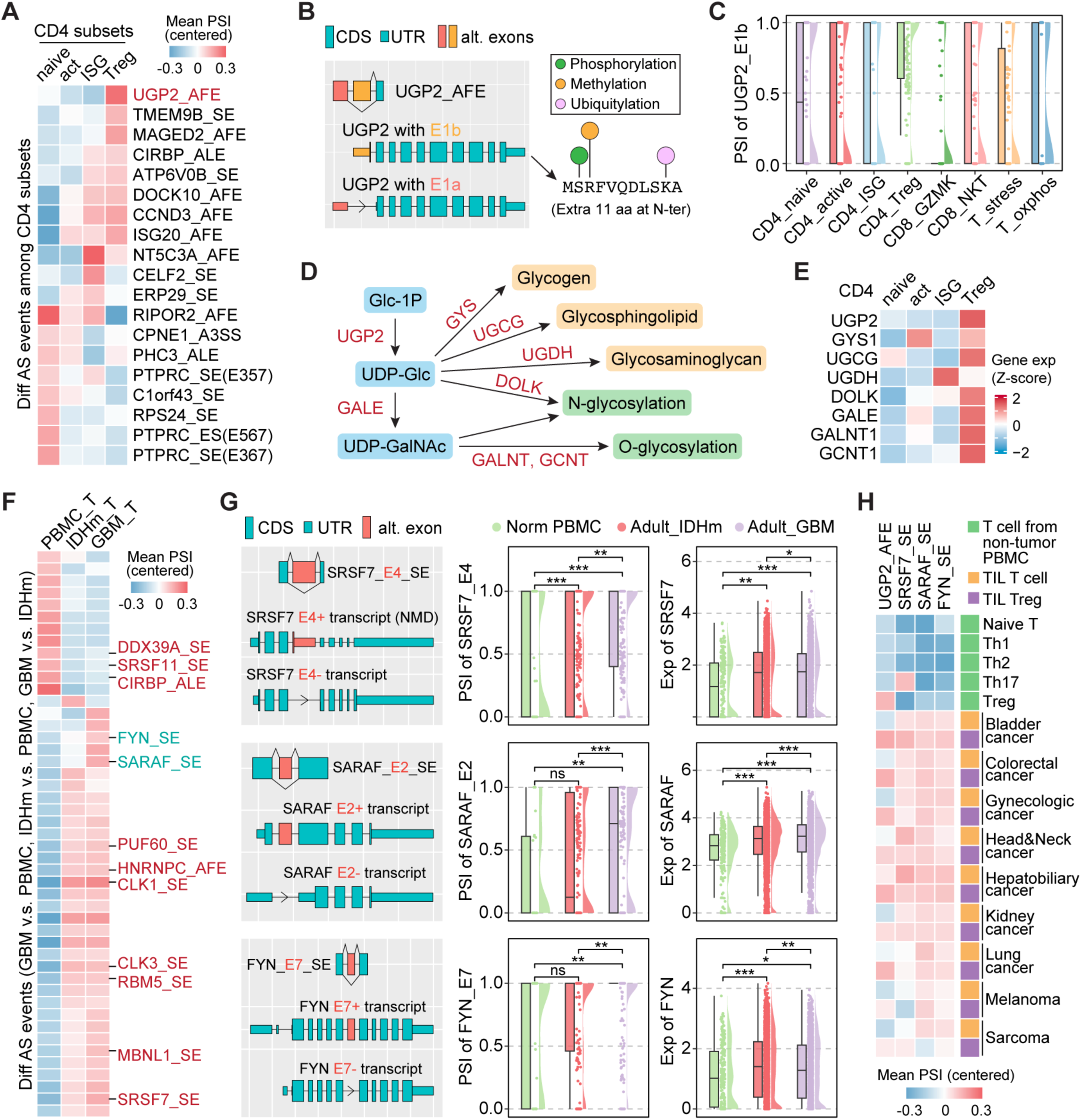
AS regulation in glioma-infiltrating T cells affects genes involved in nucleotide sugar metabolism and calcium entry. **A**. Heatmap showing the mean PSI of differential events among CD4^+^ T subsets. B. Diagrams illustrating the exon structures of UGP2_AFE. **C.** Raincloud plots showing the PSI distribution of UGP2_AFE. **D.** Metabolic pathways and associated enzymes centered on UDP-glucose. **E.** Heatmap showing gene expression across CD4^+^ T cell subsets. **F.** Heatmap showing the mean PSI of differential events among normal and glioma-infiltrating T cells. **G.** Diagrams illustrating the exon structures (left) and raincloud plots showing the PSI distribution (right). **H.** Heatmap showing the mean PSI of indicated events in bulk RNA-seq of T cell subsets from normal PBMCs and tumor-infiltrating T cells (TILs). *, p < 0.05. **, p < 0.01. ***, p < 0.001.

Since T cell subsets exhibit limited AS heterogeneity, we combined tumor-infiltrating T cells to compare their AS landscape with that of T cells from PBMCs of non-tumor patients. We identified 47 AS events that were differentially spliced among GBM-infiltrating T cells, IDH-mut glioma-infiltrating T cells, and normal PBMC T cells (Fig. 5F and Table S11). Interestingly, these events are enriched for genes involved in RNA splicing regulation (Fig. S5D). For example, glioma-infiltrating T cells exhibit increased inclusion of E4 in *SRSF7* (Fig. 5G), a critical splicing factor recently reported to promote the type I IFN response in macrophages^65^. *SRSF7*-E4 inclusion results in an NMD transcript, thereby reducing protein production^66^ (Fig. S5E).

In addition, several events distinguish GBM-infiltrating T cells from those in IDH-mut gliomas (Fig. 5F-G). One notable example involves *SARAF*, which regulates store-operated calcium entry and prevents calcium overload (Fig. S5F)^67^. Maintaining calcium homeostasis is essential for proper T cell receptor (TCR) activation^68^. In GBM-infiltrating T cells, *SARAF*-E2 is more frequently included, producing the full-length protein. In contrast, E2-excluded isoform, enriched in non-tumor PBMC T cells, lacks the entire ER luminal terminal, potentially disrupting calcium balance in T cells (Fig. 5G and S5G). Another alternative exon that is more included in GBM T cells is the E7 of *FYN* (Fig. 5G), a key kinase that associates with TCR complex and facilitates downstream signaling pathways essential for T cell activation^69^. *FYN*-E7 encodes a portion of the SH2 and SH1 domains, including the ATP-binding site, which is essential for its kinase activity^70^.

To determine whether this AS regulation in T cells is glioma-specific or a pan-cancer phenomenon, we analyzed published bulk RNA-seq data in tumor-infiltrating T cells across various tumor types and normal PBMC-derived T cells^71,72^ (Table S5). Consistently, UGP2-E1b was specifically upregulated in all Treg populations isolated from different cancers, as well as in Tregs from normal PBMCs compared to other T cell subsets (Fig. 5H and S5H). Additionally, AS events in *SRSF7*, *SARAF*, and *FYN* exhibited significantly increased PSI in tumor-infiltrating T cells compared to normal controls. In summary, these four events reflect a pan-cancer T cell regulation observed beyond gliomas.

### Glioma-associated myeloid cells include a hypoxic subset that may drive immune suppression and a remodeling subset that may support tumor growth

Lastly, we focused on the myeloid cells, the dominant immune population in the glioma TME, including both brain-resident microglia (Mg) and monocyte-derived macrophages (Mφ)^13^. Unsupervised clustering analysis identified eight clusters, with c7 highly expressing glioma-associated genes (*PTPRZ1*, *EGFR*) and c5 expressing the dendritic cell marker *ITGAX* (Fig. S6A-B). Therefore, we removed c5 and c7 and performed a de novo clustering analysis, which identified seven refined clusters (Fig. 6A). Based on canonical Mg/Mφ markers (Mg: *TMEM119*, *P2RY12*; Mφ: *CD163*, *TGFBI*), differentially expressed genes, and curated gene signatures^59,73^ (Fig. 6B-C and Table S12), these clusters were defined as follows: monocyte (*S100A12*, *FCN1*), Mφ with a hypoxia signature (Mφ_hypoxia), Mg with M1 (pro-inflammatory) or M2 (anti-inflammatory) signature (Mg_M1 or Mg_M2), mixed Mφ/Mg with an antigen-presentation signature (M_APC), mixed Mφ/Mg with an extracellular matrix remodeling signature (M_remodel), and mixed Mφ/Mg expressing stress-related heat-shock proteins (M_stress). Mg clusters dominate in IDH-mut glioma and the peripheral region of GBM, while Mφ_hypoxia and M_APC are enriched in GBM core (Fig. 6D). Deconvolution of bulk glioma data confirmed the enrichment of Mφ_hypoxia in adult IDH-wt gliomas, particularly in the necrotic regions, which correlates with a worse prognosis (Fig. 6E-F).

**Figure 6.**
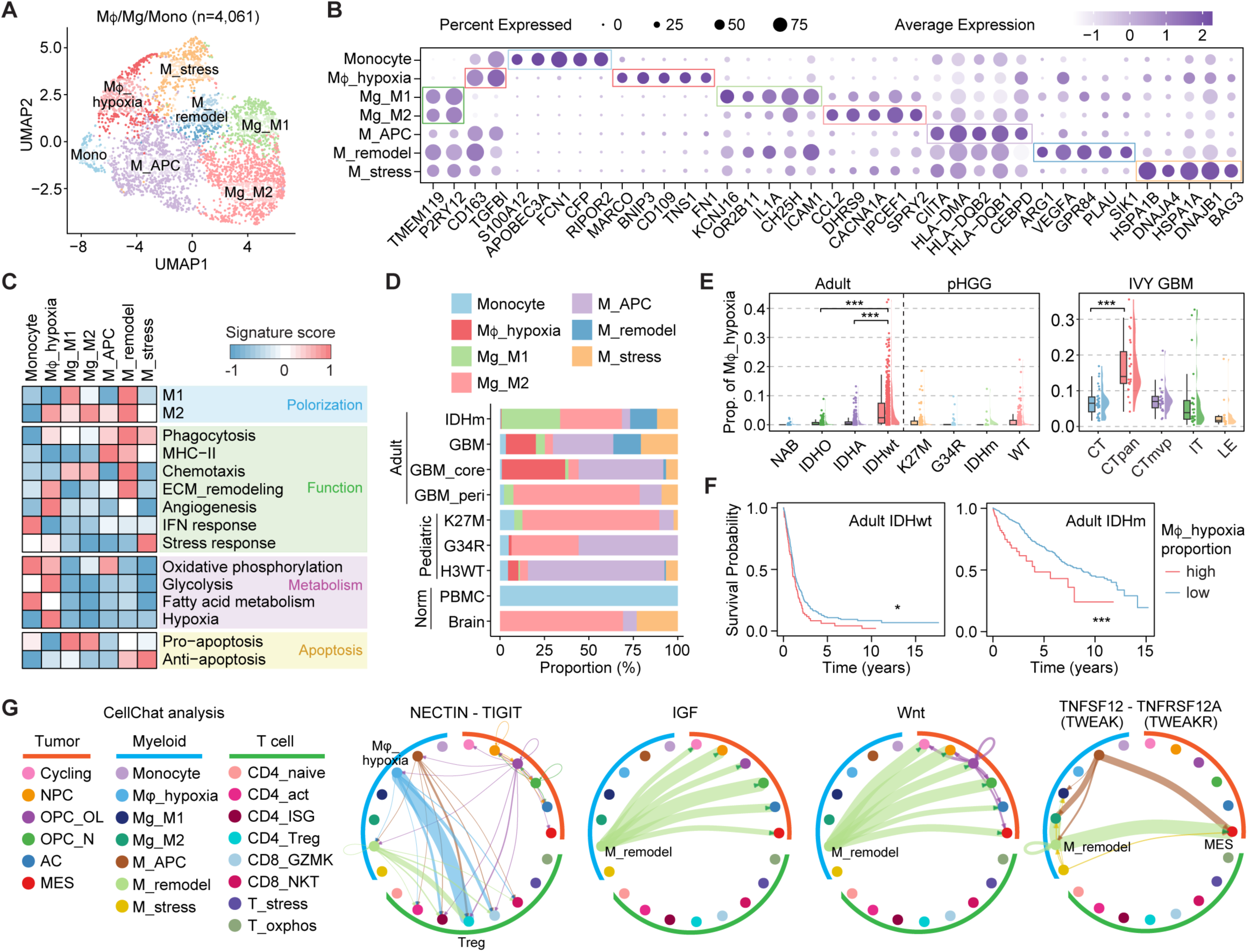
Glioma-associated myeloid cells include a hypoxic subset that may drive immune suppression and a remodeling subset that may support tumor growth. **A**. UMAP projections of glioma-associated myeloid cells colored by assigned cellular state. **B.** Dot plot showing marker gene expression. **C.** Heatmap showing scaled gene signature scores of curated gene signatures across myeloid subsets. **D.** Bar graph summarizing the compositions of myeloid cell subsets from indicated groups. **E.** Raincloud plot showing the proportion of Mφ_hypoxia subset in indicated groups, as predicted by deconvolution analysis. CT, Cellular Tumor. CTpan, pseudopalisading cells around necrosis. CTmvp, microvascular proliferation. IT, infiltrating tumor. LE, leading edge. **F.** Kaplan-Meier analyses of overall survival across adult glioma subtypes, stratified by the predicted proportion of Mφ_hypoxia. The cutoff of Mφ_hypoxia proportion is 0.1 (IDHwt) or 0.02 (IDHm). **G.** CellChat circle plot showing specific ligand-receptor interactions. The boldness of edges indicates the strength of cell-cell communication pathways. *, p < 0.05. ***, p < 0.001.

We further investigated potential cell-cell communication among glioma, myeloid, and T cell subpopulations using CellChat^74^. Notably, we identified NECTIN-TIGIT signaling crosstalk between Mφ_hypoxia and CD4_Treg (Fig. 6G). T cell Immunoreceptor with Ig and ITIM domains (TIGIT), an immune checkpoint receptor, suppresses T cell activation and proliferation upon binding to its ligands including nectin cell adhesion molecule (NECTIN) and poliovirus receptor (PVR/CD155)^75^. Additionally, we observed that M_remodel serves as a key source of ligands that activate insulin-like growth factor (IGF) and Wnt signaling pathways in glioma cells—both crucial for tumor growth and stem cell maintenance^76,77^. Furthermore, M_remodel also activates TNFRSF12A signaling in MES tumor cells, which was reported to trigger NF-κB activation and enhance glioma cell invasion^78^. In turn, MES glioma cells potentially modulate myeloid and T cell function through macrophage colony-stimulating factor (CSF) and PVR-TIGIT signaling, respectively (Fig. S6C), further shaping the immunosuppressive TME. In summary, this analysis highlights the distinct yet cooperative roles of the hypoxic and remodeling myeloid populations in glioma, collectively fostering a tumor-promoting and immune-suppressive microenvironment.

### Single-cell AS analysis in glioma-infiltrated myeloid cells reveals distinct AS patterns between microglia and monocyte-derived macrophages

A total of 580 AS events were identified as variable events in glioma-infiltrating myeloid cels (PSI-SD > 0.2, Table S13) and used for AS-based clustering analysis (Fig. 7A). Notably, the Monocyte and Mφ_hypoxia populations were distant from two Mg subsets in the UMAP plot, suggesting the AS divergence between brain-resident microglia and blood-borne macrophages. This is further supported by the largest number of differentially spliced events observed between Mφ_hypoxia and Mg_M2 (Fig. 7B-C and Table S14). Similar to T cells, a large proportion of the differential events in myeloid cells are AFE (Fig. 7B), suggesting a regulatory role of alternative transcription initiation in immune cells.

**Figure 7.**
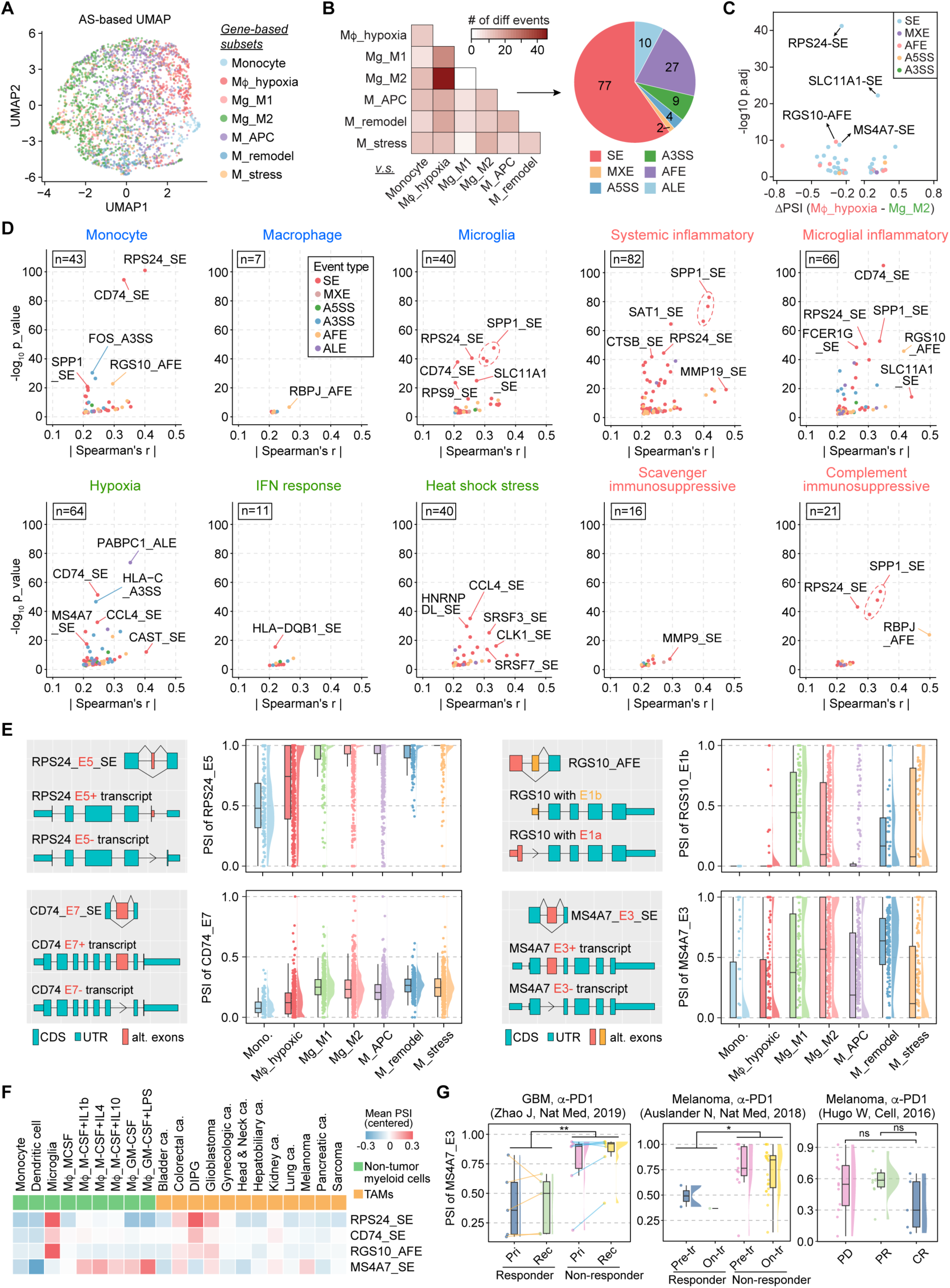
Single-cell AS analysis in glioma-infiltrated myeloid cells reveals distinct AS patterns between microglia and monocyte-derived macrophages. **A**. UMAP projections of glioma-associated myeloid cells based on the AS landscapes. Cells are colored by gene-based cellular state. **B.** Left, heatmap showing the number of differentially splicing events for each pairwise comparison of myeloid subsets. Right, number of events in each AS category. **C.** Volcano plot showing differential events between Mφ_hypoxia and Mg_M2. **D.** Scatter plot showing the log_10_ p-value (y-axis) and absolute value of spearman’s r (x-axis) for events associated with each myeloid cell program. **E.** Diagrams illustrating the exon structures of the indicated AS events. Raincloud plots showing the PSI distribution. **F.** Heatmap showing the mean PSI of indicated events in bulk RNA-seq of myeloid cells from normal PBMCs and tumor-infiltrating macrophages (TAMs). DIPG, diffuse intrinsic pontine glioma. **G.** Raincloud plot showing the PSI of *MS4A7*_E3_SE in bulk RNA-seq data of GBM or melanomas treated with anti-PD-1 therapy. Pri, primary. Rec, recurrent. Pre-tr, pre-treatment. On-tr, on-treatment. PD, progressed disease. PR, partial response. CR, complete response. The lines connect samples from the same patient. *, p < 0.05. **, p < 0.01. ns, not significant.

A recent study from Dr. Bradley Bernstein’s lab systematically investigated the immunomodulatory phenotypes in glioma-infiltrating myeloid cells by decomposing single-cell transcriptomes into discrete expression programs^79^. They disentangled cell identity programs, such as monocyte, macrophage and microglia, from cell activity programs, which include four immunomodulatory programs as well as cellular programs related to responses to hypoxia, interferon, and unfolded proteins. We performed a correlation analysis between event PSI and program usage percentage to identify AS/AFE/ALE events associated with each myeloid program (Fig. 7D and Table S15). Several events recurrently appeared across multiple programs and were strongly associated with the monocyte/microglia lineage (Fig. 7E and S7A-B). These include SE events in the ribosomal protein RPS24, where the E5+ Mg isoform, lacking the acetylation site at the C-terminus, shows increased protein stability and promotes cell survival under hypoxic conditions^80^; CD74, a chaperone protein essential for MHC-II antigen presentation, where E7 encodes a thyroglobulin type I protease inhibitor domain^81^ and is more included in Mg subsets compared with monocytes; and an AFE event in *RGS10*, which encodes a GTPase-activating protein that negatively regulates the NF-κB pathway in microglia^82^, whereas the Mg isoform of RGS10 lacks 16 amino acids at the N-terminus, containing two serine phosphorylation sites. The differential isoform usage of these three events between microglia lineage and other myeloid lineages was also observed in non-tumor myeloid cells based on our analysis of published bulk RNA-seq data^83–86^ (Fig. 7F and S7C). Analysis of pan-cancer tumor-associated macrophages (TAMs)^71,87^ showed that the Mg isoforms of these three genes are expressed in diffuse intrinsic pontine glioma (DIPG, a subtype of pHGG), adult GBM, and, surprisingly, some of the colorectal cancers (Fig. 7F).

A SE event of *MS4A7*-E3 correlates with the hypoxia program (Fig. 7D), showing the lowest PSI in monocyte and Mφ_hypoxia subsets and the highest in M_remodel subset (Fig. 7E). *MS4A7* belongs to the membrane-spanning 4A subfamily, characterized by four transmembrane domains, and its E3 encodes the first two transmembrane domains (Fig. S7B). In non-tumor myeloid cells, the PSI of *MS4A7*-E3 is highest in φM stimulated with GM-CSF plus LPS (Fig. 7F and S7C). We further examined the PSI of *MS4A7*_E3_SE event in bulk glioma RNA-seq data, as its gene expression is specific to the myeloid population (Fig. S7D). Interestingly, its PSI was associated with anti-PD1 therapy responsiveness in GBM patients, with the responder group showing a lower PSI value than the non-responder group (Fig. 7G). A similar correlation was observed in melanoma patients with anti-PD1 therapy^88,89^. Taken together, these findings highlight the potential role of AS in shaping myeloid cell identity and function within the glioma microenvironment.

## Discussion

Tumors, including gliomas, comprise a heterogeneous mix of cells in different states, with transcriptional programs shaped by their cell-of-origin lineage, genetic alterations, and microenvironmental cues. This cellular diversity underpins the complexity of tumor biology and the challenge of developing effective therapeutic strategies. Since most mammalian genes undergo AS, often generating protein isoforms with distinct functional properties, understanding the full scope of tumor heterogeneity necessitates analyzing not only gene expression but also isoform-level changes. Here, we enhance current single-cell transcriptome analysis in gliomas by incorporating the overlooked layer of AS regulation and identified multiple AS events that are differentially spliced among tumor cells, tumor-infiltrating T cells, and myeloid cells. Isoform sequence analysis and structural predictions suggest that some of these events may confer isoform-specific functions, potentially regulating tumor phenotype or immune response. Further investigation is needed to validate the functional relevance of these AS event, thereby facilitating the identification of AS-associated therapeutic targets or clinical biomarkers.

Our analysis in tumor population confirms previous findings that the MES state, enriched around the hypoxic necrotic areas of GBM, is more frequent in recurrent tumors than in primary ones and is associated with a worse prognosis^90^ (Fig. 1F, 1G, and 1J). Previous studies have identified C/EBPβ and STAT3 are key transcription factors that prevent neural differentiation and trigger reprogramming towards a mesenchymal lineage in glioma^91^. Our data highlight the potential role of another transcription factor, TCF12, in this process. TCF12 is highly expressed in undifferentiated mesenchymal stem cells, and its downregulation promotes osteoblast differentiation^92^. In gliomas, TCF12 has been reported to exert opposing functions by forming different dimers: the TCF12-TWIST dimer promotes glioma cell invasion by upregulating periostin (POSTN) expression^93^, whereas the TCF12-TCF4 dimer has the opposite effect, suppressing POSTN expression and tumorigenesis^94^. The E15-encoding region lies within an activation domain that mediates protein-protein interactions. We hypothesize that AS of TCF12-E15 alters its heterodimerization, thereby influencing the transcription of lineage-associated genes.

Cytoskeleton genes are preferentially regulated by AS during neurogenesis^95^. Our analysis comparing core and peripheral GBM cells revealed AS alterations in multiple genes involved in actin filament-organization. Notably, the TPM1 and ACTN4 isoforms expressed in peripheral GBM cells both contain unique amino acid sequences that introduce cysteine oxidation sites, potentially enabling a redox-sensitive mechanism for actin disassembly and turnover. Considering the brain’s high oxygen consumption and lipid-rich content^96^, this AS-redox co-regulation of actin dynamics may play important roles in peripheral GBM cells, particularly in their invasion and cell-cell interactions with other cells in the brain. Another shared feature of the peripheral isoforms of these genes is their ability to promote stress fiber formation in cancer cell lines^97,98^. Further investigation is needed to determine the isoform-specific function of these events in infiltrating GBM cells.

AS plays a crucial role in regulating immune cell differentiation and function^99–101^. One prominent example is the AS of CD45 in T cells, producing up to eight isoforms that characterize the transition from naive to memory states and modulate the TCR activation threshold^102^. Our analysis also detected this splicing change in glioma-infiltrating T cells (Fig. 5B). Similarly, the AS of *FYN*-E7 represents another isoform-mediated mechanism of TCR regulation^103^, which we found to be enriched in GBM-infiltrating T cells. Moreover, widespread AS alterations in splicing factor genes between glioma-infiltrating T cells and normal PBMC T cells suggest a global shift in splicing machinery. Specifically, we identified increased inclusion of a “poison exon” (PE) in *SRSF7* in glioma-infiltrating T cells. This PE inclusion, promoted by SRSF7 itself, leads to NMD, a common autoregulatory mechanism observed in many splicing factors to maintain homeostatic expression levels through a negative feedback loop^66^. It would be interesting to investigate whether disrupting this negative feedback loop with splicing-switch oligonucleotides could enhance splicing machinery function and further amplify the anti-tumor activity of glioma associated T cells.

Lastly, the AS analysis in myeloid cells further supports the lineage-specific nature of AS, as the most significant differences were observed between Mg and Mono/Mφ. It is also notable that the exclusion of *MS4A7*-E3 correlates with increased responsiveness to anti-PD-1 therapy in both GBM and melanoma patients. This E3-skipped isoform of *MS4A7* has recently been shown to promote M2 macrophage polarization in the GBM microenvironment^104^, potentially contributing to immune evasion. MS4A7 has also been reported to drive NLRP3 inflammasome activation via direct physical interaction^105^.

Further investigation is needed to explore the underlying mechanisms and assess the potential biomarker value of this splicing event in larger datasets.

In summary, this study revealed AS-level transcriptomic heterogeneity of both tumor and immune cells in gliomas, underscoring the importance of isoform-level regulation in future transcriptomic analyses. While transitioning from a gene-centric to an isoform-centric approach may require additional effort, it holds the potential to provide a deeper understanding of the complex biological processes driving tumorigenesis.

### Limitations of the study

The accuracy of PSI estimation in scRNA-seq data from the SMART seq2 platform is limited by the low sequencing depth in individual cells, which may not provide sufficient reads to reliably quantify events in low-expressing genes. As a result, the differential AS events identified in this study are primarily from moderate to high-abundance genes. Additionally, complex AS events, such as co-occurring alterations across multiple exons, are challenging to interpret using short-read RNA sequencing. In the future, applying long-read scRNA-seq, which allows an entire transcript to be captured by a single read, could enable more precise isoform-level quantification rather than event-level analysis. Lastly, while we expect our main conclusions to be robust, analyzing single-cell AS in larger cohorts with more patient samples will provide further insights. Examining different regions within the same tumor to address spatial AS heterogeneity in glioma is also an important next step.

## Resource availability

### Lead contact

Requests for further information and resources should be directed to and will be fulfilled by the lead contact, Shi-Yuan Cheng.

### Materials availability

This study did not generate new unique reagents.

### Data and code availability

All the code and related data have been deposited at Zenodo (10.5281/zenodo.15048579) and is publicly available as of the date of publication. Any additional information required to reanalyze the data reported in this paper is available from the lead contact upon request.

## Acknowledgements

This work was supported by United States National Institutes of Health (NIH) grants NS115403, NS122375, NS126810, NS125318, and Lou and Jean Malnati Brain Tumor Institute at Northwestern Medicine (S.Y.C.); United States Army Medical Research Acquisition Activity W81XWH-22-1-0374 and HT9425-24-1-0573 (X.S.); NIH CA234799 (D.M.T.). S.-Y.C. is a Zell Scholar at Northwestern University. We thank Northwestern IT Research Computing Services for providing the Quest High-Performance Computing Cluster platform and NUSeq core facility for providing service of Nanopore library prep and sequencing. We thank Dr. Mario L. Suvà, and Dr. Mariella G. Filbin for providing access to the raw single-cell RNA-seq data.

## Author contributions

S.-Y.C. and X.S. conceived the project. X.S. performed all the computational analyses, M.L. and X.S. performed data collection. Manuscript writing – Original Draft, X.S.; Review & Editing, X.S., D.T., S.-Y.C., B.H., X.Y., R.W., M.W., Q.H., and D.S. Funding Acquisition, S.-Y.C. and X.S.

## STAR Methods

### Method details

#### Single-cell RNA-seq data processing

The raw scRNA-seq data from human glioma samples, normal brains, and PBMCs were downloaded from the Data Use Oversight System (DUOS), the European Genome-Phenome Archive (EGA), or NCBI GEO, with detailed source information provided in Table S1. Clinical information of glioma patients was provided in Table S2. Reads were mapped to the GRCh38 reference genome (GENCODE release 44) using HISAT2 (v2.2.1)^106^, and gene-level counts were quantified with HTSeq-count (v2.0.2)^107^. The gene-level count data were then imported into the Seurat package (v5.2.1)^108^ in R (v4.4.0) for downstream gene-level analysis. Splice junction counts were quantified using the STAR aligner (v2.7.9a)^109^ in 1-pass mode and then imported into the MARVEL package (v2.0.5)^35^ for downstream AS-level analysis. The scripts for running HISAT2, HTSeq-count, and STAR are available on Zenodo (10.5281/zenodo.15048579) under the path “pipeline/single_cell_data_processing.sh”.

#### Copy number variation (CNV) and single-nucleotide variation (SNV) analysis in single-cell RNA-seq data

CopyKAT (v1.1.0)^26^ was used to infer CNV in single cells. Cells from the same dataset were analyzed together, except for the G34R dataset, where cells were separated by patient. Clusters assigned as non-malignant based on marker gene expression—including clusters 1, 3, 5, 6, 10, 11, 13, 15, 18, 21, 22, 23, 24, 25, 27, 28, and 29 (Fig. S1A)—were used as the normal cell reference when running CopyKAT. The detailed code for running CopyKAT was deposited on Zenodo (10.5281/zenodo.15048579) under the path “pipeline/Rscript_copyKAT.R”. SNV detection in genes *IDH1*, *H3-3A* (H3.3), *H3C2*(H3.1)*, and H3C3* (H3.1) were perform with MonoVar python package^110^. The code for running MonoVar has been deposited on Zenodo (10.5281/zenodo.15048579) under the path “pipeline/MonoVar.sh”.

#### Gene expression-based clustering, sub-clustering, and differentially expressed gene analysis in scRNA-seq data

Gene-level analysis in scRNA-seq data were performed using Seurat (v5.2.1)^108^. Low-quality cells were filtered based on the number of detected genes (1,000-15,000), total RNA counts (<4,000,000), and mitochondrial gene percentage (<20%). After quality control filtering, clustering was performed on all high-quality cells using PCA for dimensionality reduction, followed by Louvain clustering based on a K-nearest neighbor (KNN) graph. Clusters were visualized using UMAP. Tumor cells were identified based on clustering, CNV inferred by copyKAT, and SNV detected in *IDH1* and *H3* genes. Macrophage and T cell populations were identified based on clustering and expression of canonical marker genes, including *CD2* and *CD3E* for T cells and *CD14* and *CSF1R* for macrophage. Sub-clustering analysis was then performed separately on tumor cells, macrophages, and T cells by re-running dimensionality reduction, clustering, and differential gene expression analysis to reveal finer-grained cellular heterogeneity. Batch effects across datasets were corrected using the Harmony package (v1.2.3)^111^ in sub-clustering analysis. Differential gene expression analysis was performed using Seurat’s FindAllMarkers function, with the Wilcoxon rank-sum test as the statistical method. The full analysis pipeline and the processed Seurat objects have been deposited on Zenodo (10.5281/zenodo.15048579) under the folder “seurat”.

#### AS-based clustering and differential splicing analysis in scRNA-seq data

AS-level analysis in scRNA-seq data were performed using MARVEL (v2.0.5)^35^. Percent Spliced In (PSI) values were quantified for AS events with following parameters: CoverageThreshold = 5 and UnevenCoverageMultiplier = 10. The resulting PSI data is deposited on Zenodo (10.5281/zenodo.15048579) under the path “MARVEL/PSI_merge.RData”. For clustering analysis, top variable events were selected based on a standard deviation (SD) of PSI greater than 0.2, with the additional criterion that events must be expressed in a sufficient number of cells (500 cells for tumor cells and 200 cells for macrophages). UMAP-based dimensionality reduction was performed based on the PSI data of variable events after Bayesian imputation of missing PSI values, with the RunPCA function in MARVEL. Differential splicing analysis between two groups of cells were performed using MARVEL’s CompareValues function, with the Wilcoxon rank-sum test as the statistical method. The cutoff for identifying differential events was a PSI difference greater than 0.2 and an FDR-adjusted p-value < 0.1. The full analysis pipeline and processed files, including the list of all variable and differential events, have been deposited on Zenodo (10.5281/zenodo.15048579) under the folder “MARVEL”.

#### Calculation of gene signature scores and myeloid program usages in single cells

Gene signature scores for curated gene sets related to T cell or macrophage functional states^59,73^ (Supplementary Tables 9 and 12) were calculated for each cell cluster of T cells or macrophages using Seurat’s AddModuleScore function. The calculation of myeloid program usage was performed using the published method and code^79^. The following parameters were used: n_components=14, init=‘random’, update_H=False, solver=‘cd’, beta_loss=‘frobenius’, tol=0.0001, max_iter=1000, alpha=0.0, alpha_W=0.0, alpha_H=‘same’, l1_ratio=0.0, regularization=None, random_state=None, verbose=0, shuffle=False. The program usage scores were normalized to 100% for each cell. The results of program usage calculation have been deposited on Zenodo (10.5281/zenodo.15048579) under the path “other/ myeloid_program_usage.RData”.

#### Cell-cell communication analysis in scRNA-seq data

Normalized expression data of tumor cells, macrophage, and T cells from Seurat were used for cell-cell communication analysis with the CellChat package (v2.1.2)^74^. The CellChatDB.human database was used to assess signaling interactions between cell types. The full code has been deposited on Zenodo (10.5281/zenodo.15048579) under the path “pipeline/Rscript_cellchat.R”.

#### Bulk RNA-seq data processing

The raw RNA-seq data from human glioma samples, normal T cells and myeloid cells, pan-cancer tumor-infiltrating T cells and myeloid cells, and melanoma samples with anti-PD1 therapy were obtained from TCGA, CGGA, CPTAC, IVY GBM atlas project, GLASS, St. Jude cloud, EGA, NCBI Bioproject, or GEO, with detailed source information provided in Table S5. Reads were mapped to the GRCh38 reference genome (GENCODE release 44) using HISAT2 (v2.2.1)^106^. Gene-level counts were quantified with HTSeq-count (v2.0.2)^107^ and normalized using DESeq2 (v1.46.0)^112^. Splice junction counts were quantified using the STAR aligner (v2.7.9a)^109^ in 1-pass mode and then imported into the MARVEL package (v2.0.5)^35^ for calculating PSI for AFE and ALE events. rMATS (v4.3.0)^113^ was used to calculate the PSI value for SE, MXE, A5SS, A3SS, and RI events. The rMATS analysis was performed in JC mode, with a modified code to compute PSI only when more than 10 reads covered the junctions of a given event. This code has been deposited on Zenodo (10.5281/zenodo.15048579) under the path “other/ rmats_adjust_psi_JC.sh”. The scripts for running HISAT2, HTSeq-count, DESeq2, rMATS, STAR, and MARVEL are available on Zenodo (10.5281/zenodo.15048579) under the path “pipeline/bulk _data_processing.sh”.

#### Deconvolution analysis of cellular composition in bulk glioma RNA-seq data

Cell type deconvolution for bulk RNA-seq data using scRNA-seq data as reference was performed using MuSiC (v1.0.0)^114^. For the deconvolution analysis in Fig. 1H, scRNA-seq data from cells representing all tumor cellular states and other non-tumor cell types was used as a reference. For the analysis in Fig. 6E, scRNA-seq data from myeloid subsets (excluding M_stress), tumor cells (grouped as one type), and all other non-tumor cell types was used as a reference. The code for running MuSiC and the processed result files have been deposited on Zenodo (10.5281/zenodo.15048579) under the folder “MuSiC”.

#### Post-translational modification (PTM) analysis

Post-translational modification (PTM) information, including phosphorylation, methylation, ubiquitylation, acetylation, and cysteine oxidation, in specific isoforms was obtained from the PhosphoSitePlus database (www.phosphosite.org) and the CysModDB database (cysmoddb.bioinfogo.org).

#### Nanopore RNA-seq of GSC1478 cells treated with PTBP1-knockdown or control

GBM patient-derived glioma stem-like cells, GSC1478, was treated with PTBP1-knockdown (KD) or control as previously reported^15^. Total RNA was isolated using a Qiagen RNeasy Mini Kit. Poly(A)-selected RNA was used for direct cDNA synthesis and sequencing library preparation with SQK-NBD114.24 kit following the Oxford Nanopore Technologies (ONT) protocol. Libraries were sequenced on PromethION P2 solo using R10.4.1 flow cells. ONT library prep and sequencing was performed at the Northwestern University NUSeq core facility. Raw sequencing data were basecalled using guppy_basecaller (v6.2.1) and aligned to the human reference genome GRCh38 using Minimap2 (v2.17-r941)^115^. Aligned reads were visualized using IGV (v2.19.1)^116^ to inspect splicing patterns. The raw sequencing data is available from the lead contact upon request. The code has been deposited on Zenodo (10.5281/zenodo.15048579) under the path “pipeline/ Nanopore_pipeline.sh”.

#### Data visualization

Figures were generated in R (v4.4.0) using the following R packages: ggplot2 (v3.5.1)^117^, ggbreak (v0.1.4)^118^, ggdist (v3.3.2)^119^, gghalves (v0.1.4), ggpubr (v0.6.0), ggtranscript (v1.0.0)^120^, scRNAtoolVis (v0.1.0), EnhancedVolcano (v1.24.0), pheatmap (v1.0.12), ComplexHeatmap (v2.22.0)^121^, survival (v3.8-3), survminer (v0.5.6). The pathway enrichment plot in Fig.3B was generated using Cytoscape (v3.10.3)^122^. The graphic abstract was created in https://BioRender.com. The codes for generating all figures have been deposited on Zenodo (10.5281/zenodo.15048579) under the folder “Figures”.

#### Quantification and Statistical Analysis

Differential gene expression analysis and differential PSI analysis between two groups were performed using the Wilcoxon rank-sum test. For survival analysis, Kaplan-Meier curves were generated, and group comparisons were conducted using the log-rank test. Correlations between two factors were assessed using Spearman’s correlation analysis. The significance thresholds defined as p < 0.05 unless stated otherwise.

---

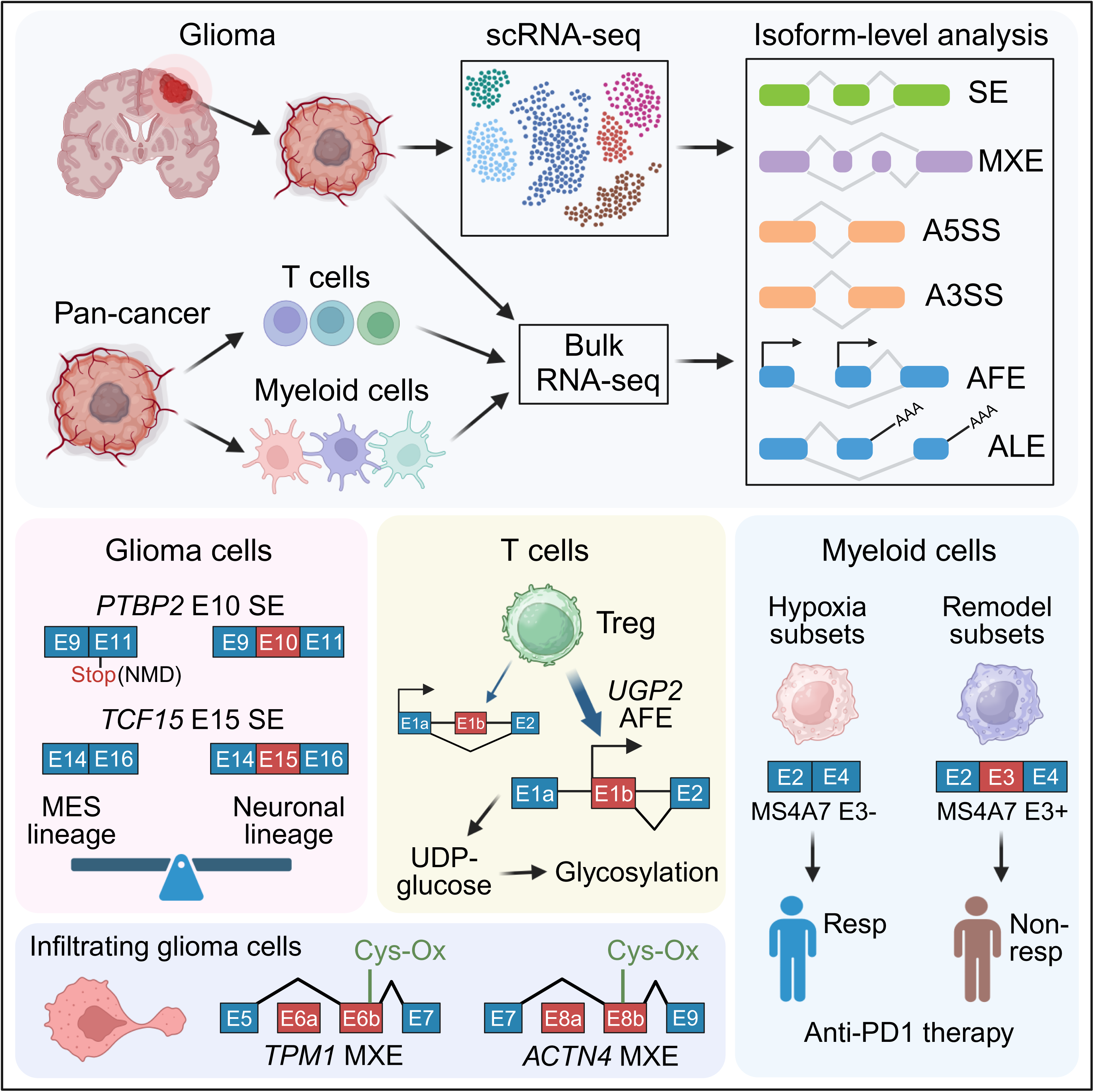

