## Supplemental Figures for "A Single-Cell Atlas of RNA Alternative Splicing in the Glioma-Immune Ecosystem"

**Document S1. Figures S1-S7**

**Document S2. Excel file containing Table S1-15**

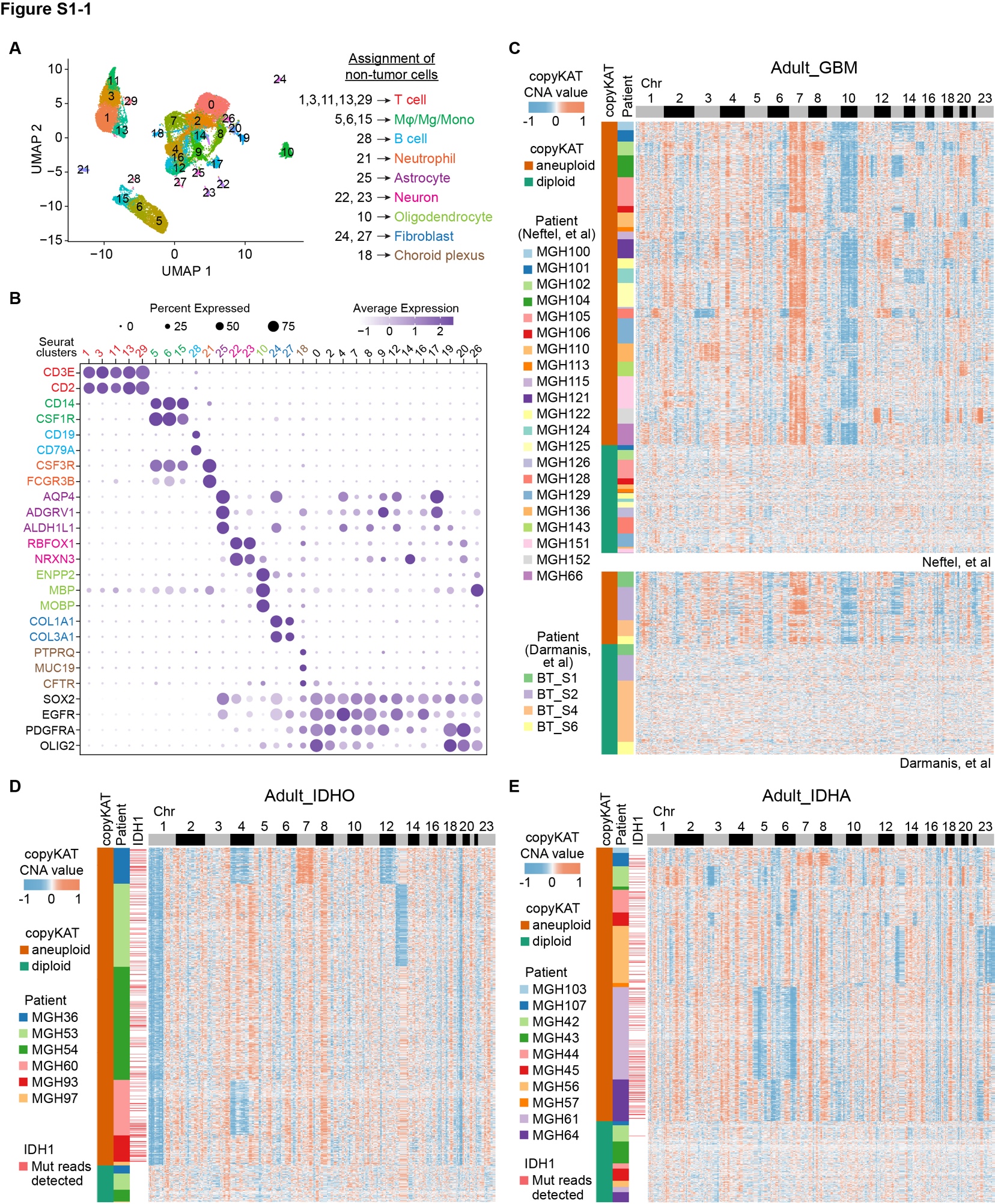

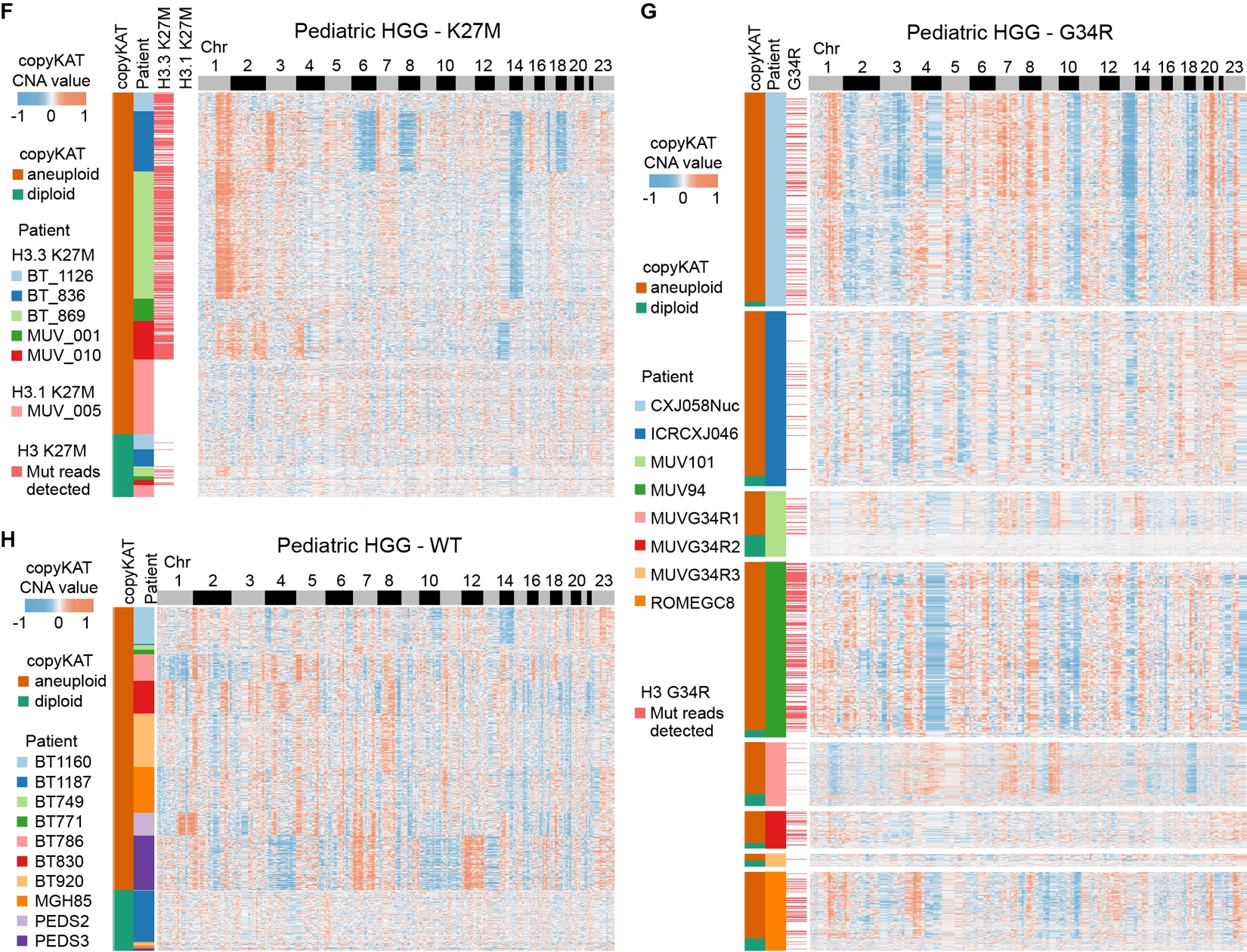

**Figure S1.** **The clustering analysis, copyKAT prediction, and SNV analysis of IDH1/H3 mutations in all high-quality single cells, related to Figure 1.**

**A.** UMAP projections of all high-quality single cells from integrated datasets of human gliomas and normal controls. Cells are colored by Seurat cluster. The assignment of non-malignant cell type was listed on the right. **B.** Dot plot showing marker gene expression for different Seurat clusters. **C-H**. Heatmaps showing the copy number variation predicted by copyKAT in indicated glioma subtypes. The annotation on the left summarizes the diploid/aneuploid status predicted by copyKAT, patient ID, and IDH1/H3 mutant reads detected from SNV analysis.

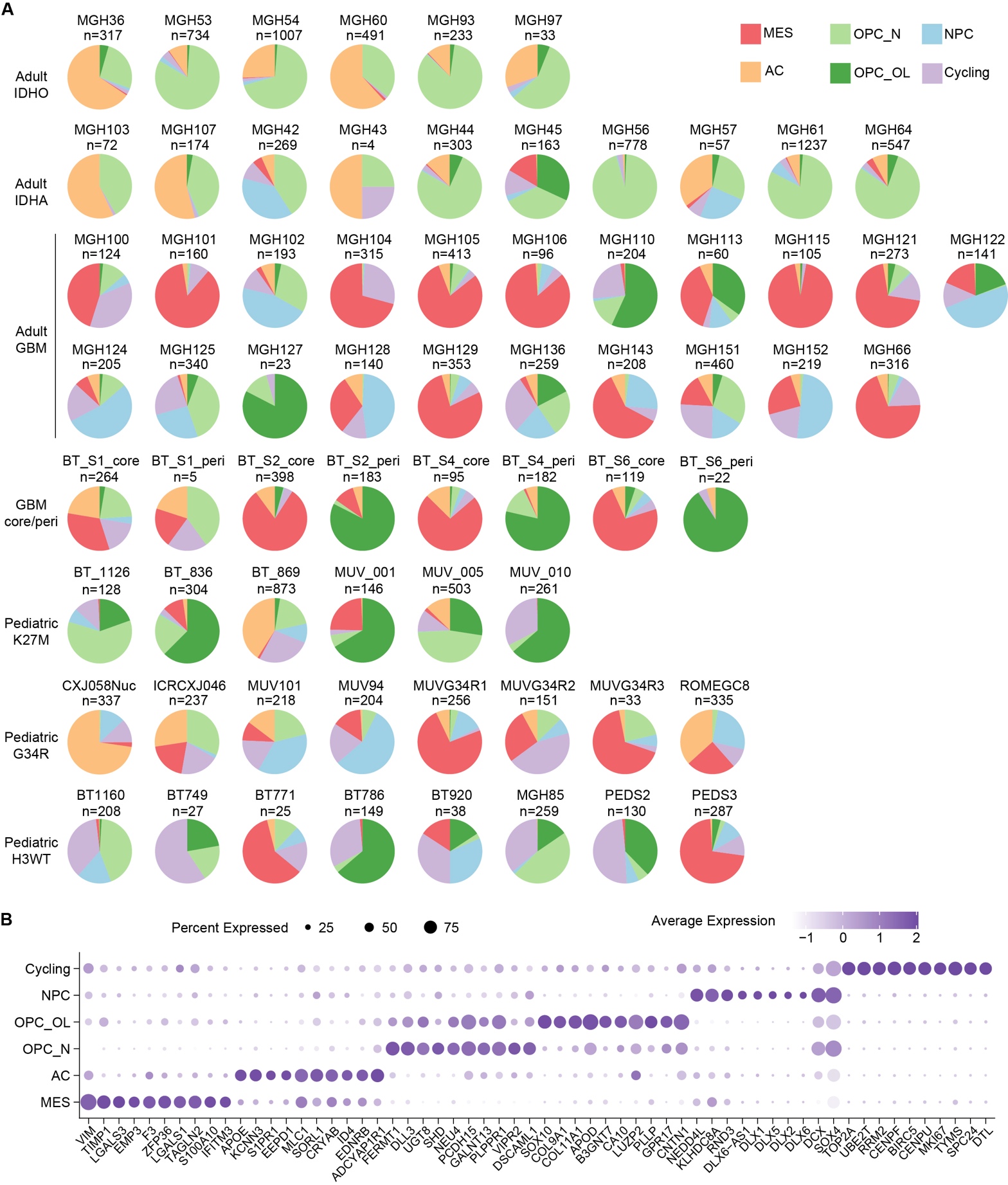
**Figure S2.** **The lineage preference of glioma cells is associated with genetic background and spatial distribution within the tumor, related to Figure 1.**

**A.** The compositions of glioma cellular states from each glioma patient. **B.** Dot plot showing the expression of top 10 marker genes for each glioma cellular state.

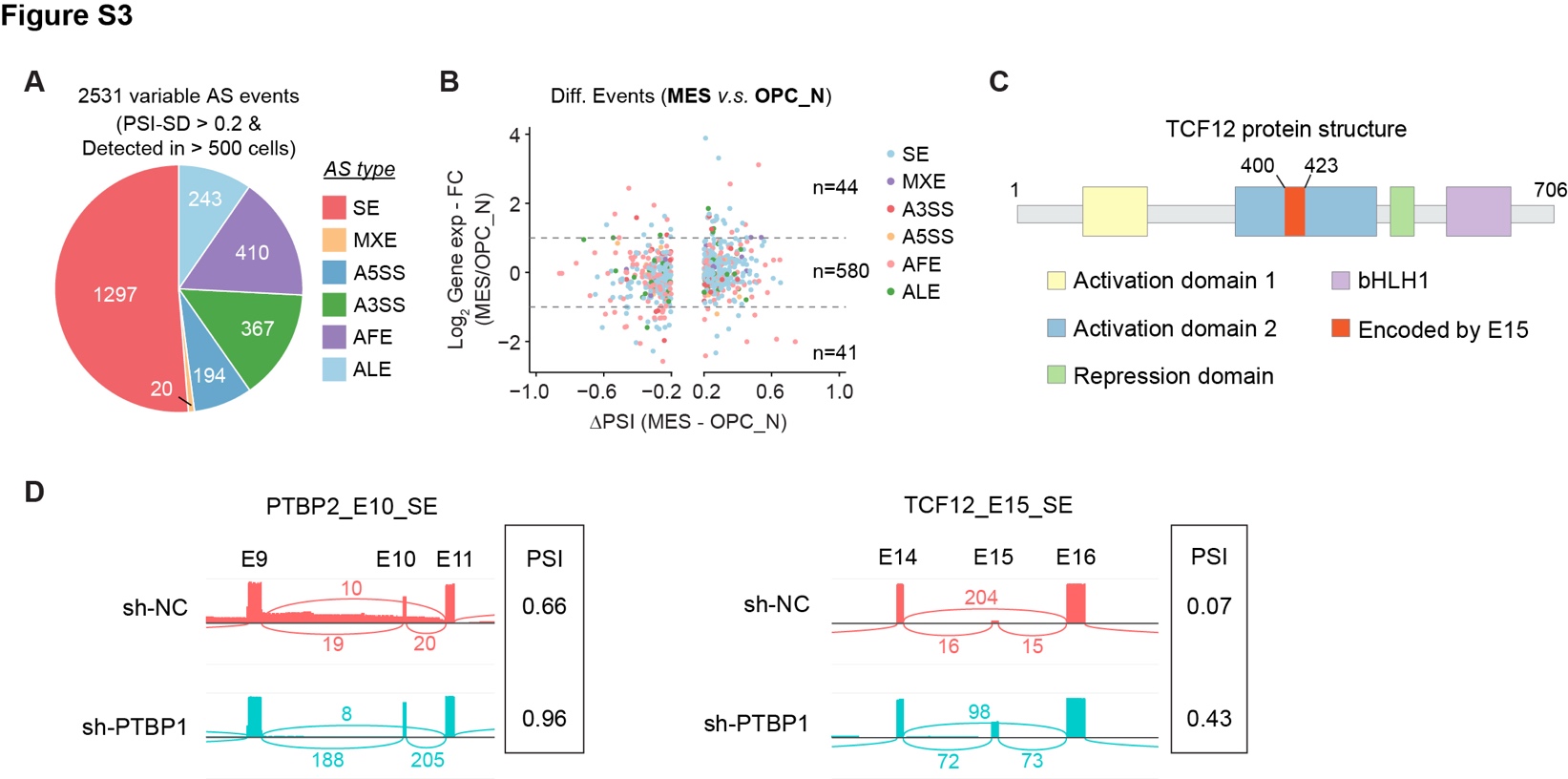

**Figure S3.** **Single-cell AS analysis in glioma tumor cells reveals lineage-specific AS regulation in *PTBP2* and *TCF12*, related to Figure 2.**

**A.** Pie graph showing the number of events in each AS category. **B.** Scatter plot showing the gene-level expression change (y axis) and PSI difference (x axis) between MES and OPC_N. **C.** Diagram showing the protein structure of TCF12, highlighting the E15-encoded region in red. **D.** Sashimi plots showing the exon usage of specific events in GSC1478 treated with shRNA targeting PTBP1 or control.

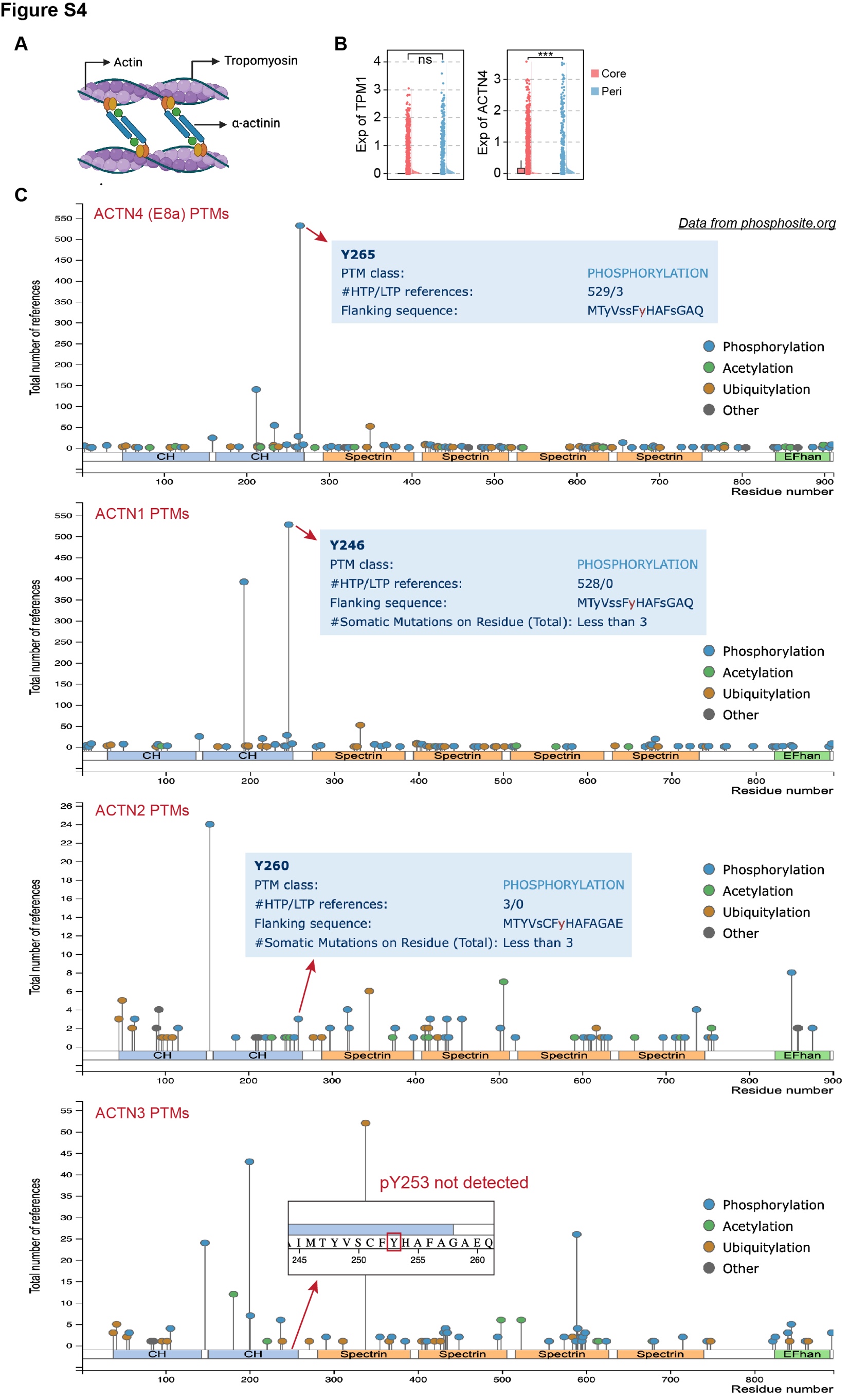

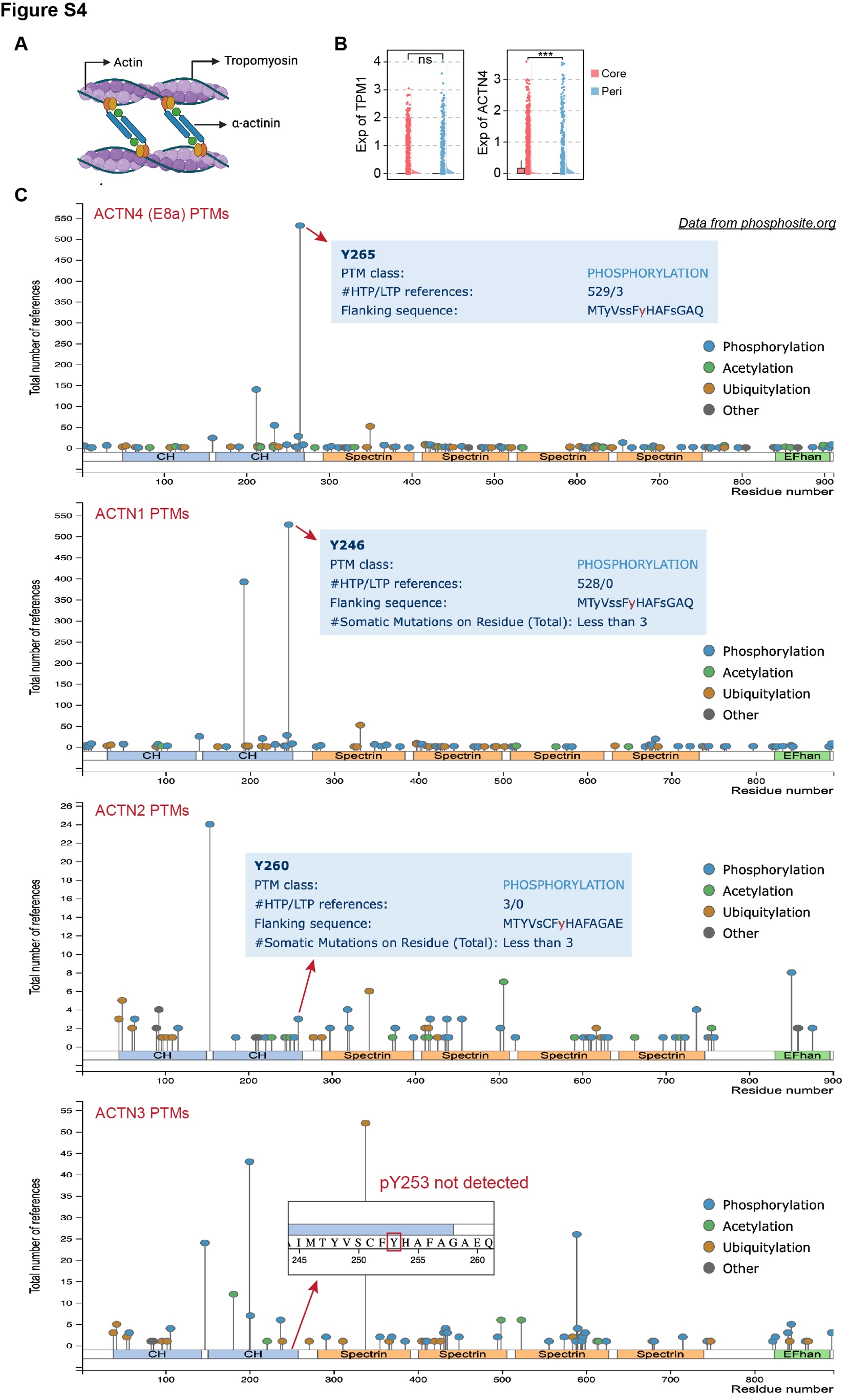

**Figure S4.** **Comparison between core and peripheral GBM cells highlights AS regulation of cytoskeleton organization, related to Figure 3.**

**A.** Schematic illustrating the roles of tropomyosin and α-actinin in actin filament organization. **B.** Raincloud plots showing the gene-level expression of *TPM1* and *ACTN4* across glioma subtypes. **D.** Reported post-translational modifications (PTMs) of four α-actinin members, sourced from PhosphoSite.org. ***, p < 0.001. ns, not significant.

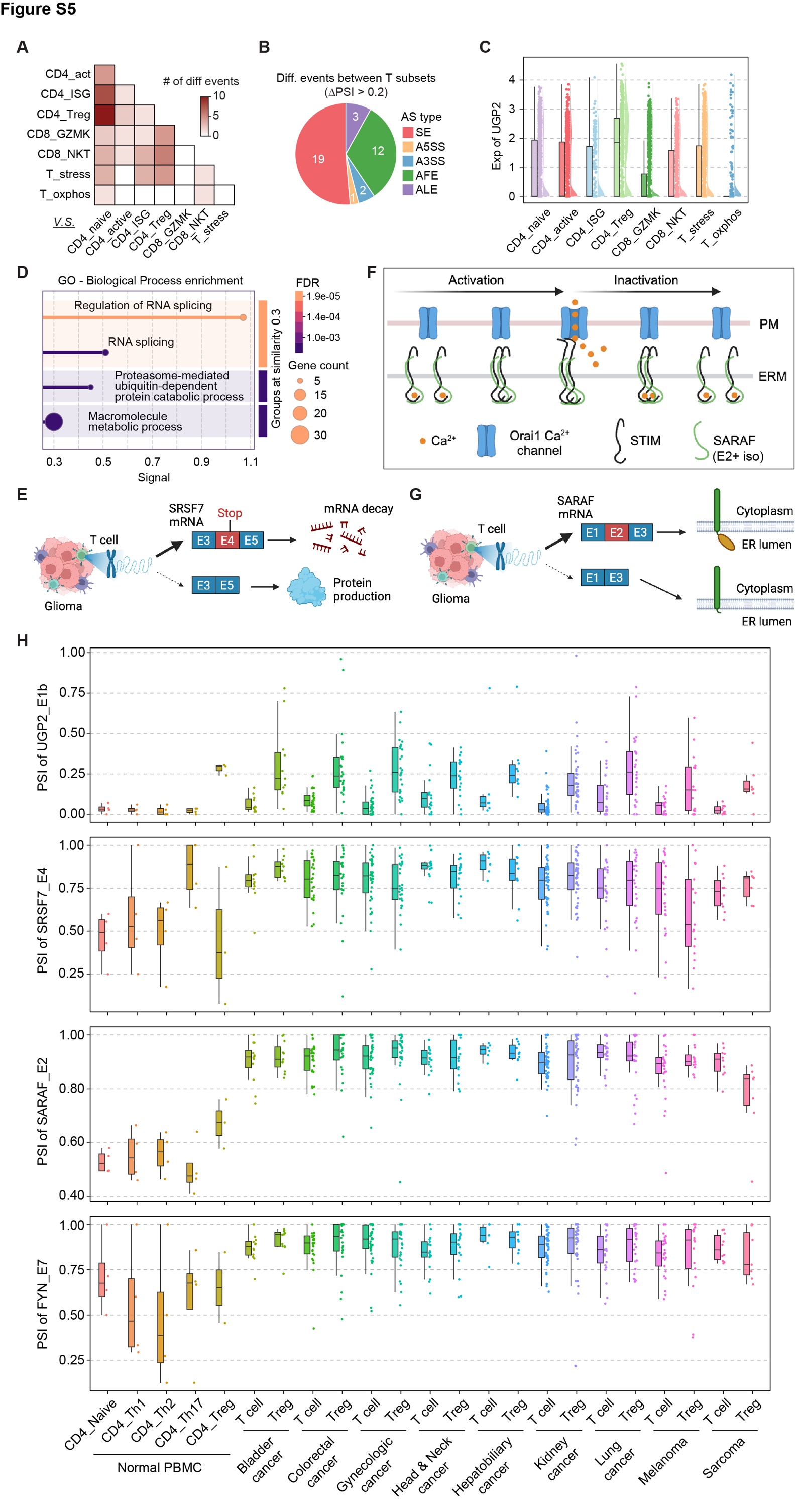

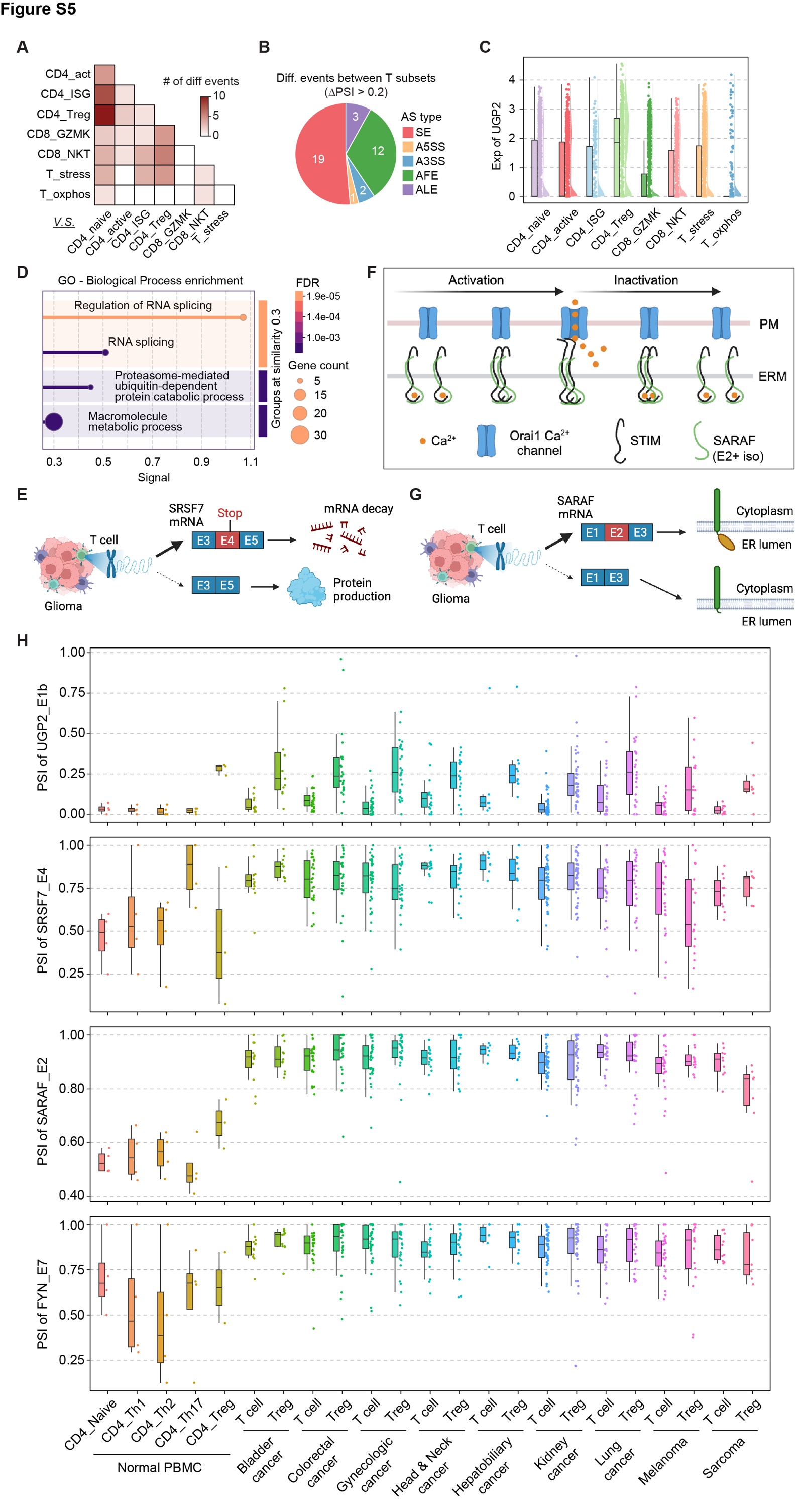

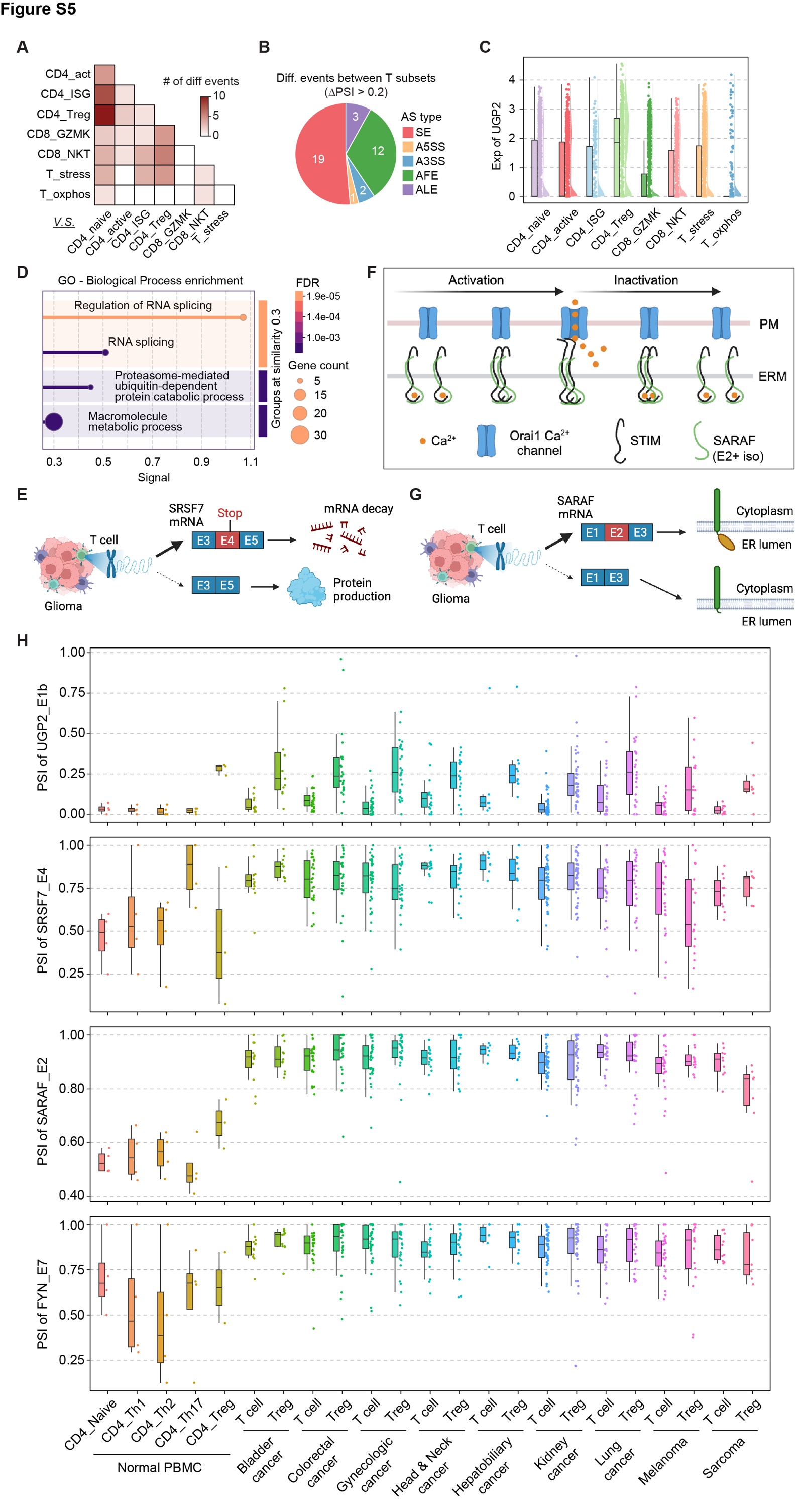

**Figure S5. AS regulation in glioma-infiltrating T cells affects genes involved in nucleotide sugar metabolism and calcium transport, related to Figure 5.**

**A.** Heatmap showing the number of differential events for each pairwise comparison among glioma-infiltrating T cell subsets. **B.** Number of differential events in each AS category. **C.** Raincloud plots showing the gene-level expression of *UGP2* across glioma-infiltrating T subsets. **D.** Enriched GO Biological Processes in genes that are differentially spliced among normal PBMC T cells and glioma-infiltrating T cells. **E.** Diagrams illustrating consequences of SRSF7_E4_SE events. SRSF7 E4+ isoform includes a pre-mature stop codon, which induces non-sense-mediated decay (NMD). **F.** Diagrams illustrating the function of SARAF in calcium entry regulation. Upon Ca^2+^ depletion from the endoplasmic reticulum (ER), the EF-hand domains of STIM proteins (located in the ER membrane, ERM) lose their bound Ca^2+^ ions. This loss triggers STIM dimerization/aggregation and induces a conformational change in their C-terminal region, enabling interaction with Orai1 channels at the plasma membrane (PM). The STIM-Orai1 interaction activates the channel, allowing Ca^2+^ influx into the cell. Once ER Ca^2+^ stores are replenished, rising Ca^2+^ levels reverse this process. In the absence of SARAF, this reversal is less efficient, leading to elevated intracellular Ca^2+^ levels. SARAF facilitates the disaggregation of STIM proteins, ensuring timely Orai1 deactivation when ER Ca^2+^ levels are restored, thereby preventing excessive Ca^2+^ accumulation in the cell. **G.** Diagrams illustrating structure difference between SARAF E2+ and E2- isoforms. **H.** Box plot showing the PSI distribution of indicated events in bulk RNA-seq of normal PBMC T cell subsets and tumor-infiltrating T cells.

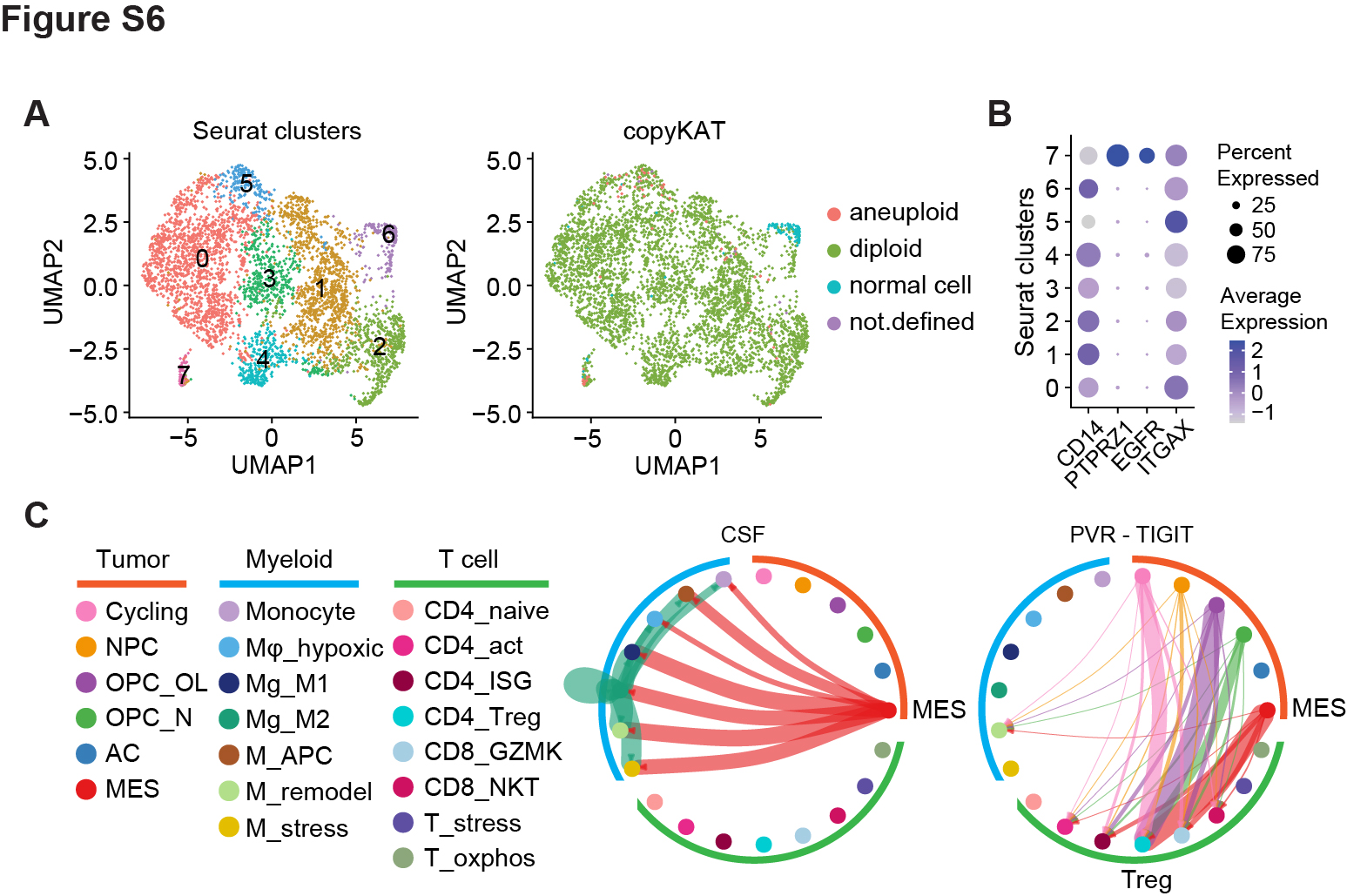

**Figure S6. Glioma-associated myeloid cells include a hypoxic population that may drive immune suppression and a remodeling subset that may support tumor growth, related to Figure 6.**

**A.** UMAP projections of glioma-associated myeloid cells colored by Seurat clusters (left) or diploid/aneuploid state predicted by copyKAT (right). **B.** Dot plot showing gene expression across Seurat clusters. **C.** CellChat circle plot showing specific ligand-receptor interactions among tumor, myeloid, and T cell subsets. The boldness of edges indicates the strength of cell-cell communication pathways.

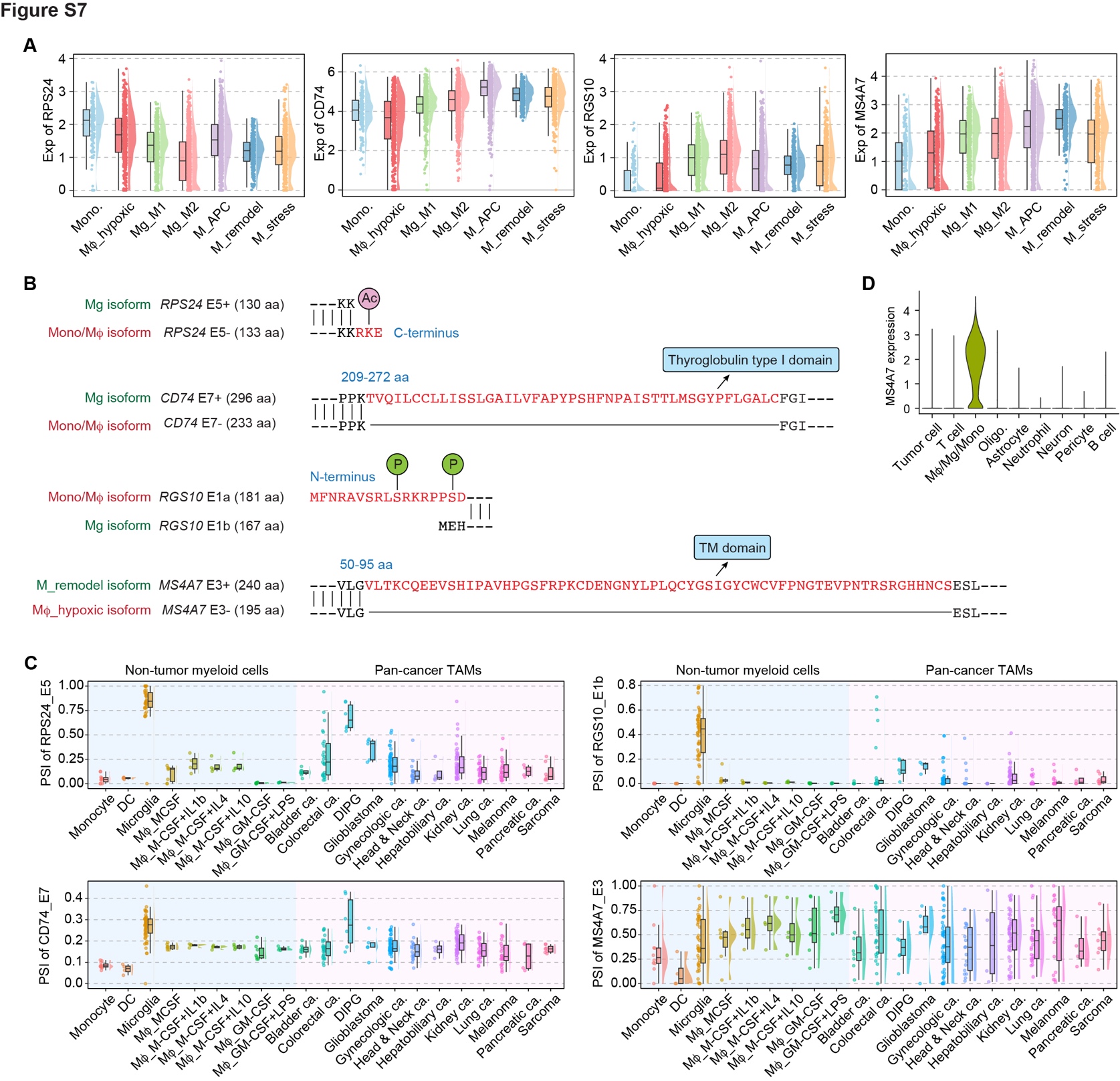

**Figure S7. Single-cell AS analysis in glioma-infiltrated myeloid cells reveals distinct AS patterns between microglia and monocyte-derived macrophages, related to Figure 7.**

**A.** Raincloud plots showing the gene-level expression of indicated genes across myeloid subsets. **B.** Differential protein sequences between indicated isoforms. The PTMs were sourced from PhosphoSite.org. **C.** Raincloud plot showing the PSI of indicated events in bulk RNA-seq data of non-tumor myeloid cells and tumor-infiltrating macrophages (TAMs) from various types of cancers. DIPG, diffuse intrinsic pontine glioma. **D.** Violin plot showing MS4A7 expression across different cell types.
